# Ventral forebrain organoids derived from individuals with schizophrenia recapitulate perturbed striatal gene expression dynamics of the donor’s brains

**DOI:** 10.1101/2022.05.26.493589

**Authors:** Tomoyo Sawada, André Barbosa, Bruno Araujo, Alejandra E. McCord, Laura D’Ignazio, Kynon J. M. Benjamin, Arthur Feltrin, Ria Arora, Anna C. Brandtjen, Joel E. Kleinman, Thomas M. Hyde, Daniel R. Weinberger, Apuā C. M. Paquola, Jennifer A. Erwin

**Affiliations:** Lieber Institute for Brain Development, Baltimore, MD, USA; Department of Neurology, Johns Hopkins School of Medicine, Baltimore, MD, USA; Department of Psychiatry & Behavioral Sciences, Johns Hopkins School of Medicine, Baltimore, MD, USA; Department of Neuroscience, Johns Hopkins School of Medicine, Baltimore, MD, USA

## Abstract

Schizophrenia (SCZ) is a brain disorder originating during neurodevelopment with complex genetic and environmental etiologies. Despite decades of clinical evidence of altered striatal function in affected patients, its cellular and molecular underpinnings remain unclear. Here, to explore neurodevelopmental alterations in the striatum associated with SCZ, we established a method for the differentiation of iPS cells into ventral forebrain organoids. Given substantial genetic heterogeneity among individuals, which can obscure disease-associated phenotypes, we generated organoids from postmortem dural fibroblast-derived iPS cells of 3 patients and 4 healthy control individuals with nonoverlapping polygenic risk score (PRS) for SCZ and whose genotype and postmortem caudate transcriptomic data were profiled in the Brainseq neurogenomics consortium. Single cell RNA sequencing (scRNA-seq) analyses of the organoids revealed differences in developmental trajectory between SCZ cases and controls in which inhibitory neurons from patients exhibited accelerated maturation. Furthermore, we found a significant overlap of genes upregulated in the inhibitory neurons in SCZ organoids with upregulated genes in postmortem caudate tissues from patients with SCZ compared with control individuals, including the donors of our iPS cell cohort. Our findings suggest that striatal neurons in the patients with SCZ carry abnormalities that originated during early brain development and a ventral forebrain striatal organoid model can recapitulate those neurodevelopmental phenotypes in a dish.

## INTRODUCTION

Schizophrenia (SCZ) affects approximately 1% of the population and often causes lifelong chronic psychosocial disturbances [1]. Both genetic and environmental factors drive the complex etiology of SCZ, which is expressed as a combination of psychotic symptoms and motivational and cognitive dysfunction [2]. The biology of SCZ is poorly understood due at least in part to heterogeneous phenotypes and the complex causal contributions of both low-penetrant common alleles and more penetrant ultra-rare variants. A recent genome-wide association study (GWAS) revealed the enrichment of common variants with genes highly expressed in both excitatory and inhibitory neurons including medium spiny neurons (MSNs) in the striatum, and no enrichment of the variants with a specific brain region [3], suggesting that a brain-wide abnormal neuronal function underlies SCZ pathogenesis. This study also demonstrated the convergence of common and rare variants associations between SCZ and neurodevelopmental disorders [3].

Neurodevelopmental abnormalities have long been proposed to play a pathogenic role in schizophrenia, a cornerstone of a neurodevelopmental hypothesis of this illness [4]. SCZ risk is thought to start accumulating *in utero* and be maintained and amplified throughout life [5]. Given the disease specificity to human and the inaccessibility of pre-disease brain tissues, studying the molecular and cellular underpinnings of SCZ has been challenging. Human induced pluripotent stem (iPS) cells and the recent progress in brain organoid technology have enabled model building to simulate aspects of human brain development. Mounting research with patient-derived iPS cells has proposed several neurodevelopmental pathways altered in SCZ, such as neural progenitor cell (NPC) proliferation, imbalanced differentiation of excitatory and inhibitory cortical neurons, in addition to altered WNT signaling during neurogenesis [6]. However, most of these studies focused on the investigation of neurodevelopmental dysfunction of cortical and/or excitatory neurons in the pathogenesis of SCZ.

Neuroimaging and neuropathological studies have demonstrated disordered connectivity between brain regions in SCZ [7-12]. One of the brain regions with disrupted function and connectivity in SCZ is the striatum [13,14]. The striatum, made up of the caudate nucleus and putamen, is a central component of the basal ganglia that form a complex neural circuit crucial for sensorimotor, cognitive, and emotional-motivational brain functions [15]. Importantly, most antipsychotics that are used for the treatment of SCZ essentially rely on the blockade of dopamine D2 receptors which are most abundant in the striatum [16]. Despite a potential central role in the pathogenesis of SCZ, cellular and molecular underpinnings of striatal dysfunction related to schizophrenia remain unclear.

The MSNs, GABAergic projection neurons carrying either D1-type and D2-type dopamine receptors, are the principal striatal neuronal population [17,18]. MSNs receive excitatory glutamatergic inputs from the cerebral cortex and the thalamus, and modulatory dopaminergic inputs from the midbrain [15]. The remaining striatal neuronal population is the aspiny interneuron, which comprises 23% of primate striatal neurons [19,20], classified into cholinergic interneurons and GABAergic interneurons. GABAergic interneurons further fall into three main subtypes expressing somatostatin (SST), parvalbumin (PV), and calretinin (CALR) [21]. During development, MSNs and interneurons originated from the subpallium, a part of the ventral forebrain. MSNs arise from progenitors in the lateral ganglionic eminence (LGE) [22-25], whereas medial ganglionic eminence (MGE) gives rise to SST- and PV-expressing GABAergic interneurons in addition to cholinergic interneurons and caudal ganglionic eminence (CGE) produces CALR-expressing GABAergic interneurons [24,26,27]. In the present study, we sought to investigate neurodevelopmental mechanisms underlying the striatal pathogenesis of SCZ. We simulated human striatum development in a dish and investigated neurodevelopmental pathophysiology underlying SCZ by generating ventral forebrain organoids (VFOs) from iPS cells.

The iPS cell-based modeling has the potential to bring about a sea change in understanding the neurodevelopmental underpinnings of SCZ, though there are important difficulties that still need to be addressed. First, SCZ is a polygenic disorder with a complex genetic architecture. Genes implicated in risk for SCZ show relatively greater expression during fetal than postnatal life [28,29] and significant overlap of GWAS risk loci between autism spectrum disorder (ASD) and SCZ [30]. However, it is unclear how potential perturbations of early brain development translates into illness in adults. Second, because there is no known morphological abnormality in fetal brains of SCZ cases, such as the increased brain size/macrocephaly in some individuals with ASD, brain organoid model validation is problematic[31-33]. Moreover, to our knowledge, the validity of the iPS cell-based SCZ models has not been evaluated with postmortem brain analysis. Although it is known that both the monolayer neuronal culture and brain organoids derived from iPS cells can recapitulate the late fetal stage of the human developing brain or preterm infant at most [34], it remains unestablished whether organoids can reproduce some disease-associated changes in the postmortem brains of adults with the diagnosis of SCZ that are possibly originated during the fetal stage.

In the present study, given these challenges in iPS cell-based modeling of SCZ, we first addressed the heterogeneity of genetic risk by analyzing VFOs derived from three individuals with SCZ and four neurotypicals by contrasting SCZ PRS of the cases with controls with nonoverlapping polygene risk scores (PRS) [35]. We also assessed the validity of the VFO model of SCZ by comparing the findings from the organoids with the data from postmortem caudate tissues including those from the iPS cell donor’s brains.

## RESULTS

### Generation and characterization of iPSC-derived ventral forebrain organoids

To investigate striatal neurodevelopmental origins of SCZ with brain organoids and explore how the iPS cell-based model can recapitulate pathophysiological changes observed in adult patient brains, we first established iPS cells from neurotypical control individuals (CT) and patients with SCZ whose genotypic and transcriptomic data of postmortem brain tissues are available through the BrainSeq consortium [36]. To maximize genetic risk differences between CT and patient, we selected four CT samples with the lowest PRS and three SCZ patients with the highest PRS among the postmortem brain cohort (Fig. 1a and Fig. S1a). We then reprogrammed postmortem dura-derived fibroblasts into iPS cells and confirmed their pluripotency (Fig. S1a, b), no genomic insertion of episomal vectors used for reprogramming (Fig. S1c), genomic integrity and genotype matching with parental brains (Fig. S1d) [37].

**Figure 1.**
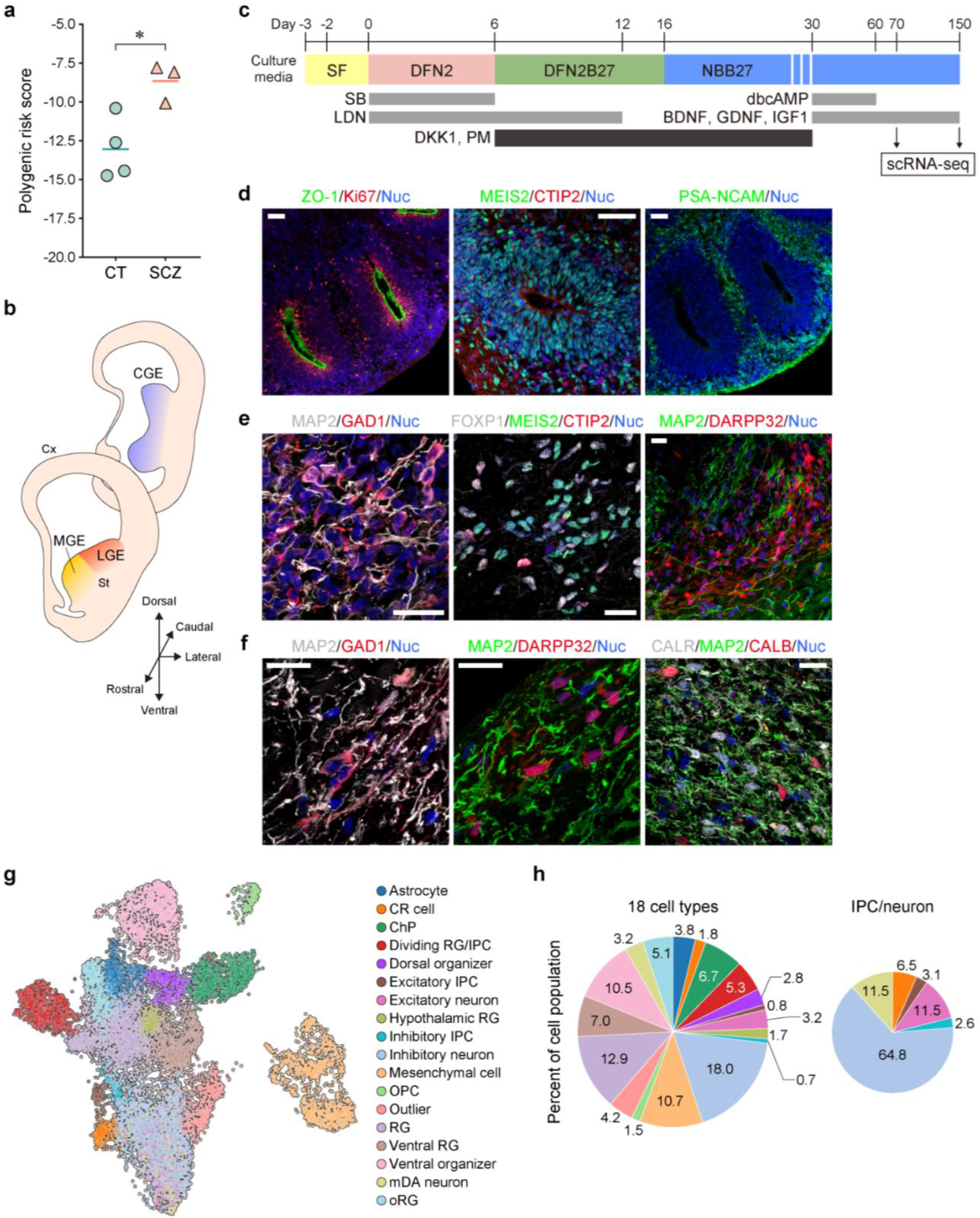
Generation and characterization of iPSC-derived ventral forebrain organoids (VFOs). **a**, Polygenic risk scores of 4 controls (CT) and 3 patients with schizophrenia (SCZ) analyzed in this study (two-tailed t-test, \**p*=0.021). **b**, Schematic of human developing brain. Cx, neocortex; St, striatum. **c**, Schematic protocol for the generation of ventral forebrain organoids (VFOs). SB, SB431542; LDN, LDN-193189; dbcAMP, dibutyryl cyclic-AMP; BDNF, brain-derived neurotrophic factor; GDNF, glial cell-derived neurotrophic factor; IGF1, insulin growth factor-1; DKK1, dickkopf related protein 1; PM, purmorphamine. **d**-**f**, Immunostaining of VFOs derived from control individual (LIBD7c6) on day 37 (**d**), day 70 (**e**) and day 150 (**f**). Nuclei are stained by Hoechst 33342 (blue). Scale bars, 50 μm (**d** and **e**), 20 μm (**f**). **g**, UMAP (uniform manifold approximation and projection) showing 15,424 cells from VFOs of 4 CT and 3 SCZ on day 70 and 150, colored by 18 major cell types. **h**, Proportion of 18 major cell types (left) and 6 major cell types among IPC/neuron population in VFOs from 7 individuals.

Given the subpallial origin of striatal neurons (Fig. 1b), we developed a protocol for differentiation of iPS cells into ventral forebrain organoids (VFOs) to simulate human striatum development by ventralizing neural cells with DKK-1, a WNT signaling pathway inhibitor, and sonic hedgehog agonist, purmorphamine (Fig. 1c). On day 12, VFOs derived from control iPS cell lines showed suppressed expression of pluripotent stem cell marker, *OCT4*/*POU5F1*, and upregulation of genes that represent induction of neural fate and differentiation of ventral and dorsal forebrain compared to undifferentiated iPS cells (Fig. S2a). On day 37, VFOs exhibited a rosette-like structure consisting of ventrally fated neuronal progenitor cells (NPCs) expressing MEIS2 and Ki67-positive proliferative cells around ventricles labeled with ZO-1. We also detected intermediate progenitor cells (IPCs)/immature neurons expressing PSA-NCAM and CTIP2 outside the structure (Fig. 1d). GABAergic neurons and cells fated to MSNs expressing FOXP1, MEIS2, CTIP2, or DARPP32 appeared in VFOs by day 70 (Fig. 1e) and we found CALR-expressing GABAergic interneurons in addition to MSNs positive for DARPP32 or Calbindin (CALB) by day 150 (Fig. 1f). To profile cellular diversity in VFOs, we performed the scRNA-seq analysis on day 70 and day 150 (n = 15,424 cells from 4 CT and 3 SCZ) (Table S1). Clustering by Leiden algorithm [61] combined with guide subclustering identified 26 clusters in VFOs, which were then annotated according to the similarity to cell clusters in cerebral organoids from a recent study [62] (Fig. S2b-c). We then further annotated the clusters into 18 major cell types (Fig. 1g) based on the expression of known marker genes (Fig. S3a, Table S2) and validated it by comparing gene expression profiles in each cell type between VFOs and the brain organoids from recent studies [38-40] (Fig. S3b-d). The 18 cell types consisted of inhibitory, excitatory, and midbrain dopaminergic (mDA) neuronal population, neuronal progenitors including proliferative cells, and non-neuronal cells, such as astrocyte and choroid plexus (Fig. 1g-h). Among IPC/neurons, 67.4% of cells were inhibitory population and the rest of 32.6 % cells were excitatory IPC/neurons (14.6%), mDA neurons (11.5%), and Cajal-Retzius (CR) cells (6.5%), indicating the ventralized neuronal lineage in VFOs (Fig. 1h). We found no unique cell type restricted to each time point (Fig. S4a, b). Thus, we decided to pool the cells from two time points together for further analyses. While VFOs on day 70 and day 150 did not significantly differ in cell type composition (Fig. S4b), differentially expressed genes (DEGs) between two time points indicated that neurons in 150-day-old VFOs were more mature than neurons in 70-day-old VFOs with upregulated synaptic genes and downregulated expression of ribosomal genes (Fig. S4c-g, Table S3).

### Inhibitory neuronal population in VFOs represented identities of immature striatal neurons

Since the present study seeks to investigate the neurodevelopmental underpinnings of altered striatal functions in the pathogenesis of SCZ, we further characterized neuronal progenitors, inhibitory IPCs, and neurons (Fig. 2a-d, Fig. S5 a-b). Neuronal progenitors including radial glia (RG), ventral RG, and outer RG (oRG) expressed marker genes for VZ in human fetal subpallium at gestational week (GW) 9-GW 12 [41], whereas inhibitory IPCs were positive for subventricular zone (SVZ) markers (Fig. 2e) and some of pre-MSN, MSN and pre-D2 type MSN (D2 MSN) markers (Fig. 2f, Fig. S5c-d, f). Inhibitory neurons rarely expressed VZ/SVZ marker genes (Fig. 2e), but a small portion of the cells was positive for immature MSN marker genes (Fig. 2f, Fig. S5c-f). Marker genes for MSN subtypes at a mature stage, such as *IKZF1, PDYN*, and *DRD1* for D1 type MSN (D1 MSN) and *EGR3, PENK, ADORA2A*, and *DRD2* for D2 MSN, were rarely detected in this cell population (Fig. 2f), suggesting that some of the inhibitory neurons in VFOs had MSN fate but they were still at an immature stage of human striatal development.

**Figure 2.**
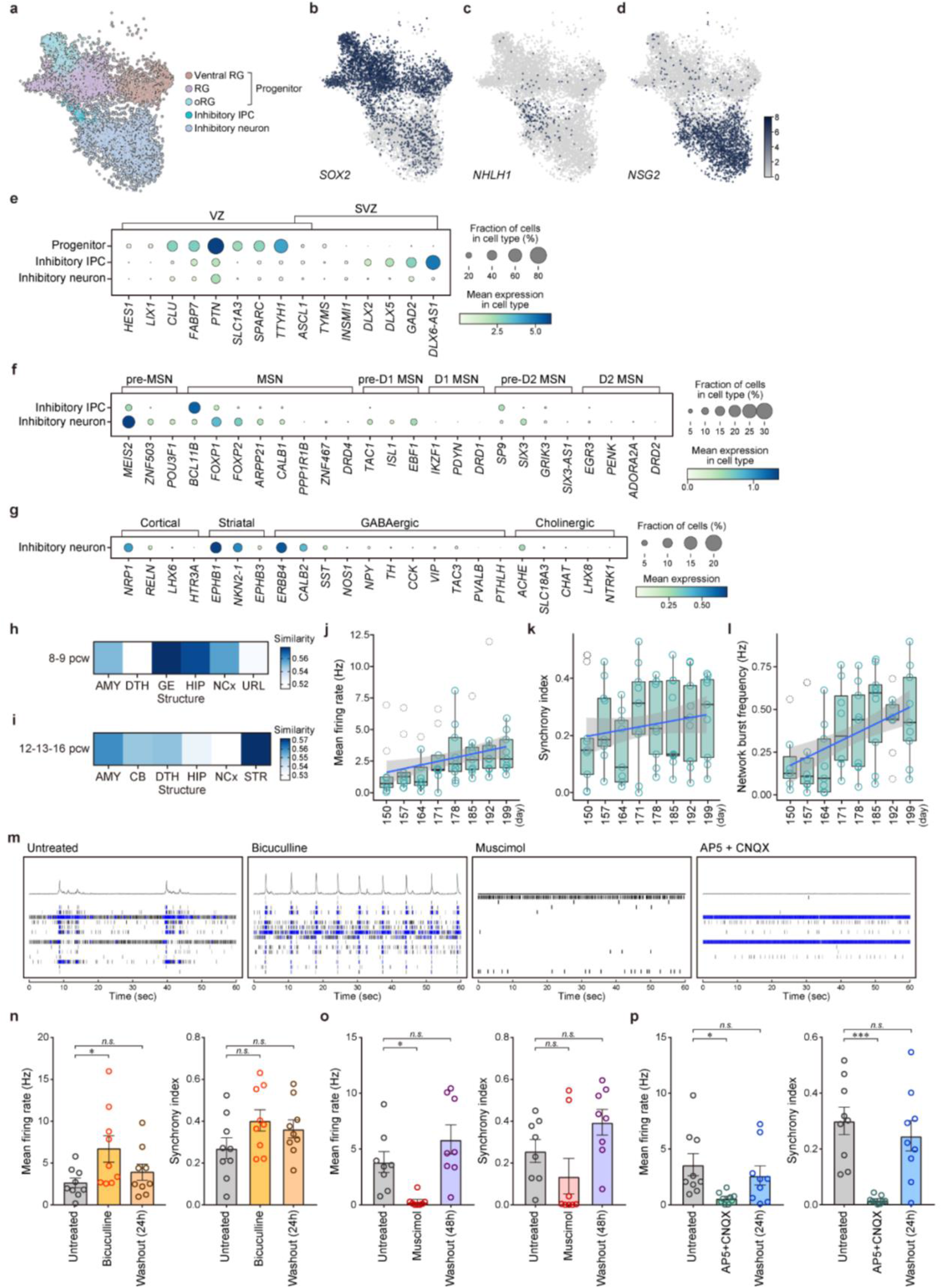
Ventral Forebrain Organoids are composed of progenitors and striatal neurons which form functional inhibitory and excitatory circuits. **a**, UMAP of neuronal progenitors and inhibitory neuronal population in VFOs (6,747 cells) from 4 CT and 3 SCZ, colored by major cell type. **b**-**d**, Representative genes expressed in progenitors (*SOX2*, **b**), inhibitory intermediate progenitor cells (IPCs) (*NHLH1*, **c**), and inhibitory neurons (*NSG2*, **d**) visualized by UMAP. Expression levels are shown in log_2_(cpm). **e**, Dot plot showing expression of selected marker genes for the ventricular zone (VZ) and subventricular zone (SVZ) in human fetal subpallium [41] and percentage of cells expressing those markers in each cell type. **f**, Dot plot showing expression of selected marker genes for medium spiny neurons (MSN) with different maturity and specific subtypes, and percentage of cells expressing those markers in each cell type among the inhibitory neuronal population. **g**, Dot plot showing expression of selected marker genes for interneuron subtypes and percentage of cells expressing those markers in inhibitory neurons in VFOs. **h**, Heatmap showing similarity of neuronal progenitors in VFOs to the BrainSpan dataset (PCW 8-9). AMY, amygdala; DTH, dorsal thalamus; GE, ganglionic eminence; HIP, hippocampus; NCx, neocortex; URL, upper (rostral) rhombic lip. **i**, Heatmap showing similarity of inhibitory IPC/neurons in VFOs to the BrainSpan dataset (PCW 12-13-16). AMY, amygdala; CB, cerebellum; DTH, dorsal thalamus; HIP, hippocampus; NCx, neocortex; STR, striatum. **j**-**l**, Time series of mean firing rate (**j**), synchrony index (**k**), and network burst frequency (**l**) of VFOs derived from a control individual (33114.c) between day 150 and day 199 (n = 3-9; VOFs for each time point). Dashed circles indicate outliers. **m**, Representative raster plots of untreated and treated control VFOs at days 220-241. **n**-**p**, Effect of treatments with bicuculline (**n**), muscimol (**o**), and AP5+CNQX (**p**) on mean firing rate and synchrony index of control VFOs (33114.c) at day 220 (**n**), day 241 (**o**) and day 227 (**p**). Data represent mean ± s.e.m. (n = 9; VOFs for recording; one-way ANOVA; ^*^*p*<0.05; ^**^*p*<0.01; ^***^*p*<0.001; n.s., not significant).

In addition to MSNs, the ventral forebrain, especially ganglionic eminences, gives rise to both striatal and cortical interneurons. We detected that more cells expressing marker genes specifying striatal interneuron (*EPHB1, NKX2-1*, and *EPHB3*) than markers for cortical interneurons (*NRP1, RELN, LHX6*, and *HTR3A*) among inhibitory neurons in VFOs (Fig. 2g, Fig. S5g). A small portion of inhibitory neurons expressed a pan-GABAergic interneuron marker, *ERBB4*, and a subtype-specific marker, *CALB2* but we found only very few cells expressing other subtype-specific markers such as *SST, NOS1, NPY, TH, CCK, VIP, TAC3, PVALB* and *PTHLH* (Fig. 2g, Fig. S5h). Given the previous studies demonstrating that majority of interneurons derived from human pluripotent stem cells do not express those marker genes at least till 20 weeks-post-differentiation [42,43], GABAergic interneurons in VFOs are still immature to express subtype-specific markers. We also found few cholinergic cells among interneurons in VFOs (Fig. 2g, Fig. S5i). Moreover, the striatum has two compartments called striosome and matrix which can be defined by their gene expression profiles and histology [44,45]. Inhibitory neurons in VFOs expressed marker genes for striosome (Fig. S5j) and matrix (Fig. S5k). To further verify the regional identity of neuronal progenitors and inhibitory neuronal populations in VFOs, we mapped the gene expression profiles of those cells onto the BrainSpan human transcriptomic dataset with the VoxHunt algorithm [46]. We found that the progenitors showed the highest similarity to ganglionic eminences (GE) when compared to human fetal brain samples at postconceptional weeks (PCW) 8-9 (Fig. 2h). In addition, inhibitory IPC/neurons exhibited the highest similarity to the striatum (STR) of human fetal brains at PCW 12-16 (Fig. 2i). We also confirmed the expression of representative marker genes for medial-, lateral- and caudal GE (MGE, LGE, and CGE) in the progenitors and inhibitory neuronal population (Fig. S6). A recent study demonstrated that the expression of *NR2F1* and *NR2F2*, which are defined markers for CGE in the rodent, was not restricted to CGE and also detected in the MGE and LGE in human fetal subpallium [41]. Given that only a few cells express other CGE-specific marker genes (*SP9, SP8*, and *CALB2*) (Fig. S6a, d), neuronal cells in VFOs are mainly composed of the MGE- and LGE lineage.

### VFOs formed functional circuits of inhibitory and excitatory neurons

To evaluate the functionality of VFOs, we performed extracellular recording of spontaneous electrical activity using multi-electrode arrays (MEAs). VFOs from a control individual were plated in a 48-well MEA plate on day 130 and recorded after day 150. We first observed a consistent increase in mean firing rate (R^2^ = 0.302, Fig. 2j) and a tendency of increases in synchrony (R^2^ = 0.163, Fig. 2k) and network burst frequency (R^2^ = N/A, Fig. 2l) along days, in addition to consistent increases in the number of spikes (R^2^ = 0.302, Fig. S7a), bursts (R^2^ = 0.445, Fig. S7b) and network bursts (R^2^ = 0.479, Fig. S7c) per minute, which indicates continuous maturation of neuronal networks in VFOs. We then examined by pharmacological intervention the role of inhibitory and excitatory synaptic transmission in forming the network (Fig. 2m). The spontaneous neuronal firing and synchronized bursting were increased by a GABAA receptor antagonist, bicuculline (Fig. 2n, Fig. S7d), and were blocked by both a GABAA receptor agonist, muscimol (Fig. 2o, Fig. S7e) and glutamate receptor antagonist (AP5; NMDA and CNQX; AMPA/kainate) (Fig. 2p, Fig. S7f). Since GABAergic neurons are known to be excitatory during early brain development and switch their mode of synaptic transmission to inhibitory upon maturation [47], we validated the expression of *SLC12A2* (also known as *NKCC1*) and *SLC12A5* (*KCC2*), which encode cation-chloride cotransporters that regulate the transition of the mode. Inhibitory neurons in VFOs expressed both of the cotransporters but the expression of *SLC12A5* was higher than *SLC12A2* suggesting that more inhibitory neurons contributed to the inhibitory synaptic transmission (Fig. S7g). Those data suggest that VFOs had intrinsic glutamatergic transmissions in which inhibitory neurons functionally modulate the neuronal activity. We also confirmed the expression of genes encoding target receptor subunits for each pharmacological intervention in VFO neurons (Fig. S7h-k). In summary, we successfully generated the VFOs enriched for functional inhibitory neurons showing immature striatal neuronal properties from human iPS cells as an *in vitro* model of human striatal development.

### Accelerated differentiation of striatal inhibitory neurons in SCZ VFOs

To understand the neurodevelopmental mechanism underlying striatal pathogenesis of SCZ, we next sought to explore cell type-specific differences in gene expression patterns between CT and SCZ. We first confirmed that the quality of cells such as total UMI counts, number of genes detected, and percentage of mitochondrial genes per cell, is not different between CT and SCZ (Table S4). Every iPS line robustly generated inhibitory neurons (>210 cells from each iPS cell line). However, the proportion of 18 major cell types varied among the iPS cell lines (Fig. S8) and between CT and SCZ (Fig. S9), consistent with previous reports demonstrating iPS cell line specification variation [48,49].

Inhibitory neurons in VFOs (Fig. 3a) from 4 CT (Fig. 3b) and 3 SCZ (Fig. 3c) showed 144 significantly differentially expressed genes, in which 45 genes were upregulated and 99 genes were downregulated in SCZ (Fig. 3d, Table S5), including upregulation of synaptic genes such as *GABRA2, NNAT* and *SCN2A* (Fig. 3e) and downregulation of genes such as *CRABP1, POU3F2* and *NOVA1* (Fig. 3f) whose dysregulation was linked to the SCZ risk [50-53]. Gene ontology (GO) analysis identified the remarkable upregulation of synaptic transmission-related genes and downregulation of genes regulating translation (Fig. 3g, Table S6). Previous studies demonstrated the global increase in translation upon neuronal differentiation and its decrease as neurons mature [54,55].

**Figure 3.**
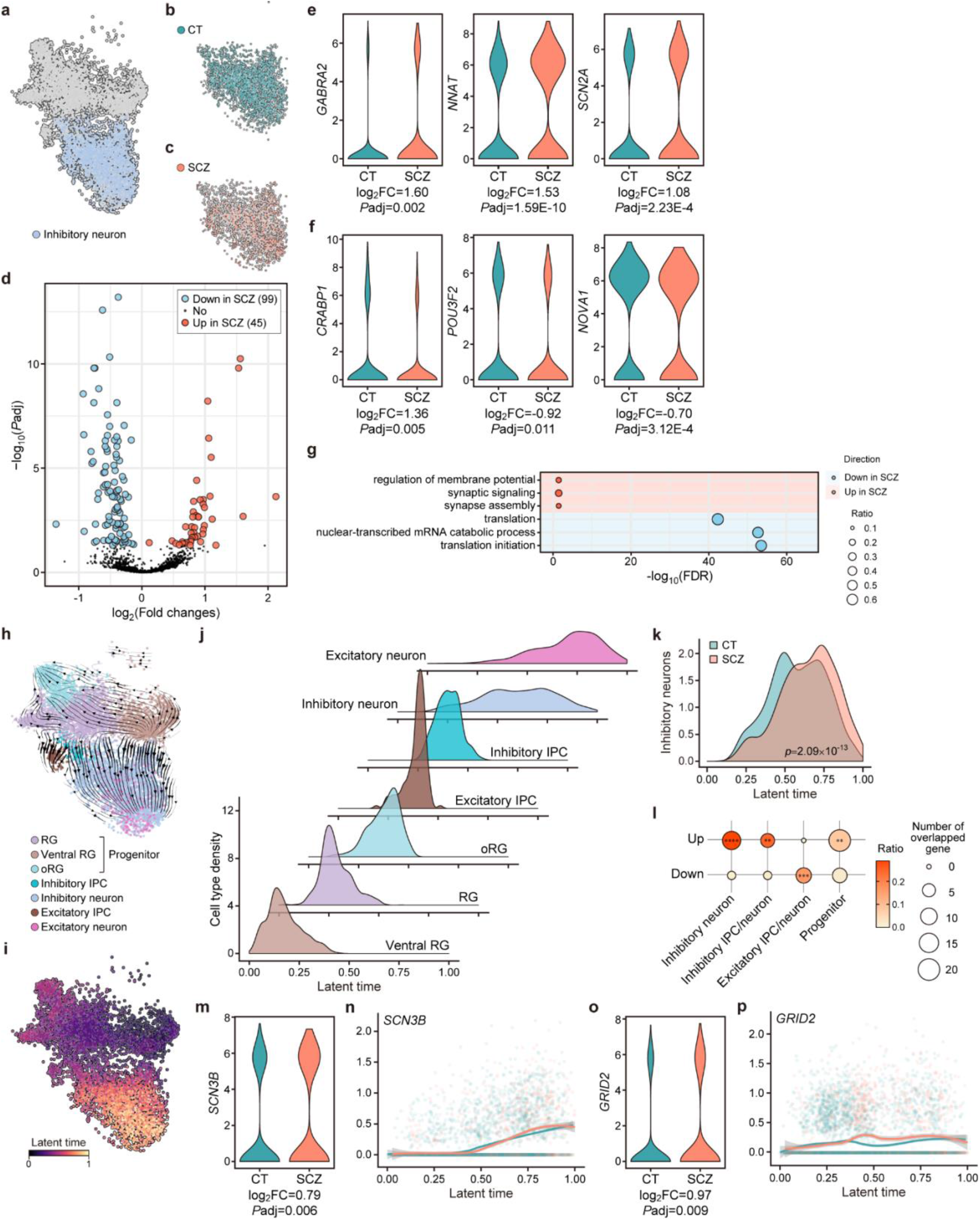
Accelerated differentiation of striatal inhibitory neurons in SCZ VFOs. **a**, Cells belonging to inhibitory neurons are colored on the UMAP of neuronal progenitors and excitatory and inhibitory IPC/neurons in VFOs from 4 CT and 3 SCZ. **b**-**c**, Cells belonging to CT (1,487 cells from 4 lines, **b**) and SCZ (1,296 cells from 3 lines, **c**) are colored on the UMAP of inhibitory neurons. **d**, Volcano plot showing differentially expressed genes (DEGs) in SCZ compared with CT. Padj, adjusted *p*-value. **e-f**, Representative genes upregulated (e) and downregulated (f) in SCZ inhibitory neurons. FC, fold change. Expression levels are shown in log_2_(cpm). **g**, Dot plot showing representative GO terms enriched for upregulated and downregulated genes in SCZ inhibitory neurons. The size of the circles indicates a ratio of genes annotated in each GO term to all 2,301 genes analyzed. FDR, false discovery rate. **h**, Velocity estimates projected onto UMAP of neuronal progenitors and excitatory and inhibitory IPC/neurons in VFOs from 4 CT and 3 SCZ. **i**, Latent time projected onto UMAP of neuronal progenitors and excitatory and inhibitory IPC/neurons in VFOs from 4 CT and 3 SCZ. **j**, Density of each of 7 major cell types applied for scVelo along latent time. k, Difference in the density of inhibitory neurons along latent time (Kolmogorov-Smirnov test). l, Dot plot showing the overlap between DEGs and 100 putative drivers for each cell type (Fisher’s exact test; **p<0.01; ***p<0.001; ****p<0.0001). The ratio of DEGs overlapped with drivers to all up- /down-regulated genes in each cell type is indicated by color. **m**-**p**, Expression of representative driver genes that were upregulated in SCZ. Expression of the genes in inhibitory neurons in CT and SCZ VFO (**m, o**). Expression levels are shown in log_2_(cpm). Expression of the genes along latent time in CT and SCZ VFOs (**n, p**).

We observed the similar enrichment of synaptic GO terms in upregulated genes and translational GO terms in downregulated genes in inhibitory IPC/neurons in DEG analysis by adding inhibitory IPCs to inhibitory neurons (Fig. S10a-e, Table S5,6). Overall, these findings suggest that inhibitory neuronal cells in VFOs from SCZ are more mature than the cells in CT VFOs.

To investigate whether the accelerated maturation of neuronal cells is specific to inhibitory lineage, we next performed DEG analysis on excitatory IPC/neurons in VFOs (Fig. S11a-c). There were 78 upregulated and 49 downregulated genes in SCZ compared to CT (Fig. S11d, Table S5), including *ANK3* (Ankyrin-3) (Fig. S11e), which is known as a common genetic risk factor for neurodevelopmental and psychiatric disorders including autism spectrum disorder [56,57], bipolar disorder [58,59] and schizophrenia [60,61]. GO analysis showed enrichment of genes contributing to translation among upregulated genes and genes associated with synaptic transmission and neurogenesis among downregulated genes (Fig. S11f), indicating decelerated maturation of excitatory neuronal cells in SCZ VFOs. We also performed DEG analysis on neuronal progenitors in VFOs (Fig. S12a-c) and identified 575 significantly differentially expressed genes in which 243 genes were upregulated and 332 genes were downregulated in SCZ compared to CT (Fig. S12d, Table S5). Interestingly, we found some of GWAS [62] significant genes among upregulated genes in SCZ, such as *CLU, PTN* and *TCF4* (Fig. S12e). GO analysis showed marked upregulation of factors associated with neurogenesis and downregulation of translational machinery in progenitors in VFOs from SCZ (Fig. S12f, Table S6). Given that the progenitors showed similarity to GE (Fig. 2h), these findings also indicated accelerated maturation of inhibitory neuronal cells. We thus demonstrated that there is a cell type-specific difference in developmental trajectory between CT and SCZ, in which the inhibitory neuronal population derived from iPS cells from SCZ exhibit accelerated maturation. These findings led us to further investigate differences in neurodevelopmental trajectory between CT and SCZ by RNA velocity analysis [63,64]. The velocity field map reflected the dynamics of neuronal differentiation as these cells transition from ventral RG or RG to neurons, with or without passing through oRG or IPC phases (Fig. 3h). The inferred latent time also demonstrated the neurodevelopmental trajectory, in which RG or ventral RG were produced earlier than oRG or IPC and neurons were produced lastly (Fig. 3i, j). We first examined cell type-specific differences in the trajectory found that excitatory IPC/neurons overall were at a later stage in the progenitor-to-neuron trajectory in VFOs compared to inhibitory IPC/neurons (Fig. S13a). However, when we look at the difference between CT and SCZ, we found that inhibitory neurons (Fig. 3k) and inhibitory IPC/neurons (Fig. S10f) had more cells at later developmental stages compared to CT cells, consistent with the results from DEG analyses suggesting the accelerated maturation of inhibitory neuronal cells in SCZ (Fig. 3g, Fig. S10e). Moreover, a delayed developmental state was found in excitatory IPC/neurons of VFOs from SCZ in comparison with CT (Fig. S11g), supporting the decelerated maturation phenotype suggested by DEG analysis (Fig. S11f). We also observed promoted developmental state of the neuronal progenitors from SCZ VFOs compared to the cells from CT VFOs (Fig. S12g), which is also indicative of accelerated maturation of inhibitory neuronal population in early brain development in the individuals with SCZ. In summary, we found the promoted developmental state of the inhibitory neuronal population in SCZ VFOs.

The RNA velocity analysis also identified putative driver genes of transcriptional changes in each cell type (Fig. S13b-e, Table S7). Driver genes display pronounced dynamic behavior and are involved in directing lineage fate decisions. Some driver genes showed overlap with SCZ risk genes identified in GWAS [62] and TWAS studies with postmortem dorsolateral prefrontal cortex (DLPFC) [65] and caudate [66] (Fig. S13f, Table S8). Notably, drivers for neuronal progenitors were significantly enriched with GWAS significant genes for SCZ (Fig. S12f, Table S8). Furthermore, genes upregulated in SCZ were significantly overlapping with the driver genes in inhibitory neurons, inhibitory IPC/neurons, and the progenitors (Fig. 3l, Table S9), whereas there was also a significant overlap between downregulated genes in SCZ excitatory IPC/neurons and their driver genes (Fig. 3l, Table S9), indicating that cell type-specific differences in the neuronal maturity arose from differences in the expression of key genes that regulate transcriptomic dynamics. We also found that some of the driver genes overlapped with SCZ-associated genes in postmortem brain of adult individuals. For example, *SCN3B* (Fig. 3m) and *GRID2* (Fig. 3o) were genes upregulated both in inhibitory IPC/neurons in SCZ VFOs and postmortem caudate from individuals with SCZ compared to CT [66]. A previous study *SCN3B*, encoding a subunit of voltage-gated sodium channels responsible for the generation and propagation of action potentials in neurons [67], showed upregulation only at the later developmental time (Fig. 3n). In contrast, *GRID2* (glutamate ionotropic receptor delta type subunit 2) began to be upregulated in SCZ cells earlier (Fig. 3p). *GRID2* was shown to be uniquely expressed in human LGE at pcw 9 among MGE, LGE and CGE [40].

We also identified *ANK3* as a driver gene for excitatory IPC/neurons, which was also downregulated in excitatory IPC/neurons in VFOs from SCZ (Fig. S11e) and overlapped with TWAS significant genes for SCZ from the analyses on postmortem caudate [66]. In the neuronal population in VFOs, *ANK3* was downregulated in SCZ cells at an immature state compared to CT and exhibited consistent and slight downregulation in SCZ throughout the latent time (Fig. 11h). Furthermore, *PTN* (Pleiotrophin), a secreted heparin-binding growth factor that regulates diverse cellular functions such as cell proliferation, neuronal migration, and neurite outgrowth [68], was a driver for the neuronal progenitors whose expression was upregulated in the progenitors in VFOs from SCZ compared to CT (Fig. S12e). *PTN* is one of GWAS [62] significant genes that was also identified as a SCZ TWAS significant gene in postmortem caudate [66]. At an earlier latent time in the progenitor-to-neuron trajectory, *PTN* was slightly downregulated in the cells from SCZ, whereas it was rapidly upregulated in SCZ along with the timing of the emergence of oRG and showed no difference in the expression level as the cells mature into neurons (Fig. S12h).

The RNA velocity analysis demonstrated the cell type-specific differences in neurodevelopmental trajectory between CT and SCZ. Inhibitory neuronal cells mature at a faster rate but the excitatory population showed decelerated maturation in SCZ VFOs compared to the cells in VFOs from CT. When we examine the proportion of cells from VFOs on day 70 and day 150 in each cell type, no significant enrichment of the cells on day 150 in SCZ VFOs was identified in the inhibitory neuronal and progenitor population (Fig. S14a-c). In addition, we observed that excitatory IPC/neurons in SCZ had more cells from VFOs on day 150 than in CT (Fig. S14d). Thus, we excluded the possibility that the differences in neurodevelopmental trajectory arise from the difference in the experimental age of the cells. In summary, our findings from cell type-specific analyses of the VFO model of human striatal development proposed the acceleration of inhibitory neuronal differentiation in striatal developmental trajectory as a molecular underpinning of striatal pathogenesis of SCZ.

### VFOs recapitulate cell type-specific transcriptional changes of postmortem brain tissues of SCZ individuals

The findings of cell type-specific perturbation of neurodevelopmental trajectory in SCZ VFOs and the identification of SCZ-risk genes in DEGs between CT and SCZ VFOs drove us to assess whether organoids can reproduce disease-associated changes in the postmortem caudate of adults, the corresponding brain region of VFO, with the diagnosis of SCZ. To address this challenge, we examined the overlap between cell type-specific DEGs in VFO model and GWAS significant genes from PGC2+CLOZUK [62], TWAS significant genes and DEGs between SCZ and CT in postmortem caudate tissues from BrainSeq study [66], which includes the donors of the iPSC cohort, and frontal cortices from PsychENCODE project [69] (Fig. 4). We did not identify a significant enrichment of the cell type-specific DEGs in VFOs with GWAS significant genes (Fig. 4a). However, we notably found a significant overlap of upregulated genes between SCZ VFOs and SCZ caudate (Fig. 4a). Downregulated genes in inhibitory neurons in SCZ VFOs were significantly depleted in upregulated genes in postmortem SCZ caudate and none of upregulated genes in other cell types showed substantial enrichment with them (Fig. 4a), suggesting a specificity of the enrichment. When we compared cell type-specific DEGs in VFOs and the postmortem cortex dataset in PsychENCODE project [69], we found a significant overlap of upregulated genes between progenitors in VFO and adult cortices (Fig. 4b), consistent with previous studies showing that the genes implicated in risk for SCZ show relatively greater expression during fetal than postnatal life [28,29]. Interestingly, although it was not statistically significant, we observed higher overlap of DEGs in excitatory IPC/neurons in VFOs and TWAS significant genes in adult cortices (Fig. 4b). In contrast, we did not identify marked overlap between DEGs in VFO inhibitory neuron and postmortem cortex dataset (Fig. 4b). These findings also indicate the cell type-specificity of the recapitulation of the disease-associated transcriptional changes with a brain organoid model. In summary, our data suggest that striatal inhibitory neurons in the patients with SCZ carry abnormalities that originated during early brain development and the VFO model can recapitulate those neurodevelopmental pathophysiological changes in a dish.

**Figure 4.**
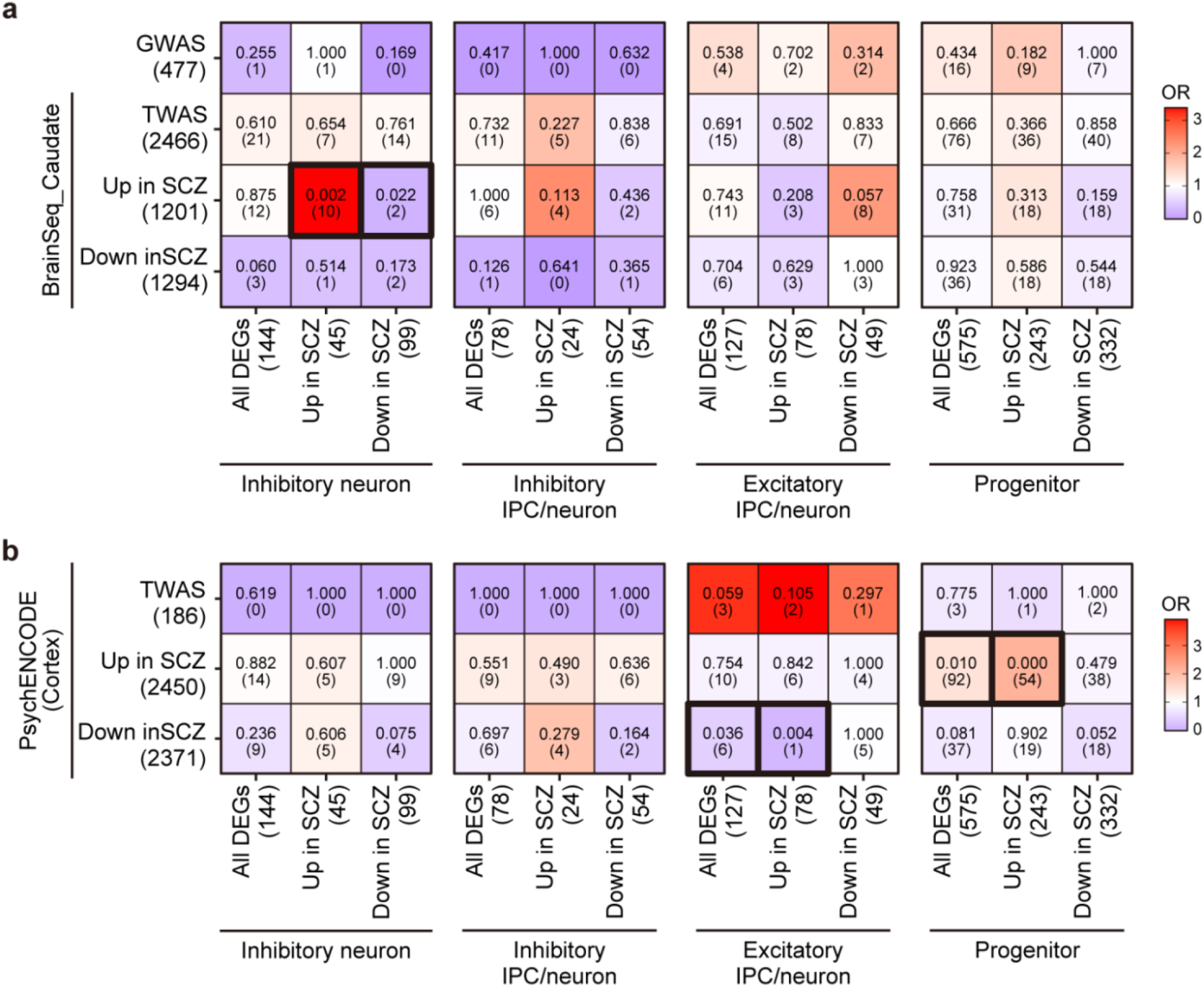
Inhibitory neurons from SCZ patients recapitulate transcriptional perturbations of SCZ postmortem caudate. Heatmap showing enrichment of cell type specific DEGs with GWAS significant genes [62] and TWAS significant genes and DEGs in postmortem caudate tissues from individuals with SCZ compared to CT [66] (**a**) and TWAS significant genes and DEGs in postmortem PFC tissues from individuals with SCZ compared to CT [69] (**b**) (Fisher’s exact test; p-value and number of genes overlapped are indicated for each comparison). OR, odds ratio. Number of genes is indicated in brackets.

## DISCUSSION

Genetic background is known as a large source of variability between iPS cell lines [70-72], which may obscure, exaggerate or mislead the disease-related phenotypes observed in the studies for disease modeling. Given the polygenic nature of SCZ and the significant overlap of the SCZ PRS between cases and controls in the general population [73], the strategy to scale back the influence of genetic variation is required for iPS cell-based modeling of SCZ, especially in the sporadic cases. Analysis of twins discordant for the disorder [74-77] and modeling the disorder with a relatively large-scale iPS cell cohort are promising strategies [78]. These strategies may, however, not be easily applicable in the majority of studies, because of the rarity of twins cases and an extremely large number of individuals theoretically required that was estimated to be necessary [71]. Therefore, in the present study, we took an alternative strategy to maximize the distance in genetic risk between patient and contrast groups. We selected three individuals with SCZ who carry the highest PRS and four neurotypical individuals with the lowest PRS among the postmortem brain donors in the LIBD brain repository (https://www.libd.org/brain-repository/). We then analyzed iPS cell-derived VFOs from the case-control cohort with significantly different PRS for SCZ.

In addition to genomic heterogeneity, it has also been a challenging endeavor to demonstrate the validity and disease relevancy of the iPS cell-based model for adult-onset psychiatric disorders, considering the maximum maturity of neural cells that can be differentiated from human iPS cells [34]. Our strategy enabled us to analyze iPS cell-derived brain organoids and postmortem brain tissues from the same individuals and thus to evaluate whether the organoid model can recapitulate any disease-relevant gene expression changes identified in the postmortem adult brain tissues.

We have demonstrated that at least at the transcription level, inhibitory neurons in VFOs are mature enough to recapitulate disease-associated changes in adult SCZ brains. We identified significant overlap of upregulated genes between inhibitory neurons in SCZ VFOs and postmortem caudate tissues of schizophrenics (Fig. 4a), implying the ability of VFOs to recapitulate pathophysiological changes observed in adult patient brains, despite the discrepancy in cellular maturity between the brain organoids and postmortem brains and the inconvenient fact that postmortem brains reflect the consequences of a life-long illness and pharmacological intervention. Though further studies are required to address these difficulties, by comparing the cell type-specific gene expression profiles in the iPS cell-derived brain organoids with postmortem caudate tissues including the same donors, the present study implied a contribution of aberrant fetal striatogenesis to SCZ pathogenesis.

Neural progenitors and inhibitory neurons in the VFOs generated in this study were at the early stage of human striatal development and had similar gene expression patterns as human GE and early fetal striatum, respectively (Fig 2h-i). Nevertheless, putative driver genes, which regulate transcriptional switches upon differentiation or maturation, for not only inhibitory IPCs/neurons and Excitatory IPC/neurons but also for neuronal progenitors showed notable overlap with genes differentially expressed in the postmortem caudate tissues between SCZ and CT [66] (Fig. 3l, Table S9). These findings suggest that at least some of the transcriptional changes in postmortem brains of patients with schizophrenia have early neurodevelopmental origins and/or maintain the consequences of *in utero* neurodevelopmental perturbations throughout life. Cell type-specific profiling of gene expression in VFOs revealed that the alterations in the developmental trajectory of inhibitory neurons begin at the progenitor stage (Fig. S12g). Several studies have demonstrated the importance of SCZ GWAS risk genes in brain development [28-29]. Interestingly, the driver genes of neural progenitors showed significant overlap with GWAS risk genes for SCZ [62], whereas the drivers for IPC/neurons did not (Fig. S13f, Table S8), suggesting that SCZ risk starts accumulating *in utero* and is coded by genomic sequences involved in striatal pathogenesis.

Furthermore, our findings of cell type-specific alterations in neurodevelopmental trajectory in VFOs, in which differentiation of inhibitory neurons is accelerated whereas it is delayed in excitatory neurons, implicate an excitation and inhibition (E-I) imbalance during early brain development in SCZ. E/I balance is essential for the formation of an optimal neural circuit, which in turn controls complex behaviors by supporting information processing. Cumulative evidence [79-82] including our recent study with cerebral organoids from monozygotic twins discordant for psychosis [74] has linked E-I imbalance to the pathogenesis of neurodevelopmental and psychiatric disorders, particularly ASD and SCZ. Moreover, recent studies identified ASD-like behavior deficits [83] and abnormal brain network dynamics [84] caused by E-I imbalance in the cellular and mouse models of neurodevelopmental disorders. Additionally, Kang and colleagues demonstrated that Disrupted-in-schizophrenia 1 (DISC1), a risk gene for major mental disorders including SCZ, leads to E-I imbalance in mature dentate granule neurons [85]. Therefore, further study to explore how the excitatory/inhibitory imbalance of developmental trajectory affects the formation and maturation of functional neural circuits is also required for understanding the neurodevelopmental etiology of SCZ.

In this study, we generated the VFOs from iPS cells to simulate human striatal development in a dish and identified cell type-specific alterations in neurodevelopmental trajectory as a molecular underpinning of striatal pathogenesis of SCZ with the iPS cell cohort consisting of three patients carrying high PRS of SCZ and four neurotypicals with low PRS. To our knowledge, this is the first study that evaluated the validity of the iPS cell-based brain organoid model of SCZ with its corresponding postmortem brain tissues from the same donors. Given the limited number of case and control individuals analyzed in the iPS cell cohort, however, the present study may still not be able to reduce undesired effects of genomic variation between individuals adequately. Also, the striatal neurons in our organoid model were at an immature stage and the consequences of altered striatal development in the pathogenesis of SCZ remain to be elucidated. Although the organoids can simulate human brain development more appropriately than 2D neuronal cultures, it is still a simplified model compared to in vivo development. Simulating developmental trajectory combined with genetic perturbations and integrating multi-dimensional organoid and brain tissues datasets would have a great impact on predicting the consequences of neurodevelopmental perturbations that are encoded by genomic risks in the patient brains. Further study with our strategy of analyzing the brain organoids and corresponding postmortem brain tissues from the same individuals with a larger case-control cohort in combination with the tracing of human brain development *in silico* could advance our understanding of the neurodevelopmental origin of SCZ and accelerate the search for the novel therapeutic target of the disorder.

## MATERIALS AND METHODS

### Tissue collection

Postmortem dural tissues of four neurotypical Caucasian males [37] and three Caucasian males with schizophrenia were collected at autopsy [65,66,86] and fibroblasts were isolated from dura mater as previously described [87]. Participants were selected from the LIBD brain repository (https://www.libd.org/brain-repository/) based on the polygenic risk score (PRS) of schizophrenia [88,89] and fibroblast availability, and were gender and race matched (Fig. 1a, Fig. S1a). PRSs are a measure of cumulative genomic risk [89] calculated as the sum of risk alleles of index single nucleotide polymorphisms (SNPs) from GWASs on schizophrenia [88,90], weighted for the strength of association, that is, the Odds Ratio [88]. Consistent with the standard procedure for PRS calculation [89], only autosomal SNPs were included in the analysis, to prevent bias related to sex. As previously described [88-91], we performed linkage disequilibrium pruning and “clumping” of the SNPs, discarding variants within 500 kb of, and in r2 ≥ 0.1 with, another (more significant) marker. We multiplied the natural log of the odds ratios of each index SNP by the imputation probability for the corresponding effective allele for the Odds Ratio at each variant, and we summed the products over all variants, so that each subject had whole genome PRSs. Ten PRSs (PRS1–10) were calculated using subsets of SNPs selected according to the GWAS P value thresholds of association with schizophrenia: 5e−08 (PRS1), 1e−06 (PRS2), 1e−04 (PRS3), 0.001 (PRS4), 0.01 (PRS5), 0.05 (PRS6), 0.1 (PRS7), 0.2 (PRS8), 0.5 (PRS9), and 1 (PRS10). PRS6 was employed to select the samples, following previous evidence [88] that it has the highest accuracy in predicting the respective disease or trait. Collection and use for research of fibroblasts were approved by the Western Institutional Review Board. Dura-derived fibroblasts were grown and maintained as previously described [37].

### Generation and maintenance of iPS cells

Dura-derived fibroblasts between passages 5 and 7 were reprogrammed with episomal vectors [92]. Plasmids pCXLE-hOCT3/4-shp53-F (Addgene #27077), pCXLE-hSK (#27078), and pCXLE-hUL (#27080) were transfected into fibroblasts using a 4D-Nucleofector system with P2 Primary Cell 4D-Nucleofector X Kit (Lonza; program DT-130). Three to five weeks after reprogramming, single colonies were picked and expanded on SNL76/7 feeder cells (ATCC, SCRC-1049) treated with mitomycin C (Sigma Aldrich) in 20% KSR medium [92]. iPS cells were then transferred to feeder-free culture on Cultrex Reduced Growth Factor Basement Membrane Matrix (Trevigen)-coated plates in StemFlex medium (Gibco). Cells were passaged every 4-6 days with Versene solution (Gibco). A control iPS cell line (MIN09i-33114.C) was obtained from WiCell and maintained in the same way [93]. All lines have undergone extensive characterization for identity, pluripotency, and exogenous reprogramming factor expression [37] (Fig. S1).

### Generation of ventral forebrain organoids (VFOs) from iPS cells

To generate VFOs, iPS cells cultured on Cultrex-coated 6-well plates in StemFlex medium were treated with 10 μM Y-27632 (BioGems) at 37°C overnight when they reach 50-60% confluency. Cells were then incubated with StemPro Accutase (Gibco) at 37°C for 7 min and dissociated into single cells. Dissociated iPS cells were plated on 96-well V-bottom plates (S-bio) at 9,000 cells/150 μL per well in StemFlex medium with 30 μM Y-27632 and incubated at 37°C for 2 days. The StemFlex medium was gradually switched to DFN2 consisting of DMEM/F12 + GlutaMAX (Gibco) supplemented with 1 × N-2 supplement (Gibco), 1 × non-essential amino acid (NEAA, Gibco), 100 μM 2-mercaptoethanol (2-ME, Gibco) and 1 × Antibiotic-Antimycotic (Gibco) from day 0. From day 6, the medium was gradually switched to DFN2B27 consisting of DMEM/F12 + GlutaMAX supplemented with 1 × N-2 supplement, 1 × NeuroCult SM1 without vitamin A (StemCell Technologies), and 1 × Antibiotic-Antimycotic. On day 12, each embryoid body (EB) was embedded in a droplet of Matrigel Basement Membrane Matrix Growth Factor Reduced (Corning) as previously described [74,94] and transferred to a 60-mm culture dish in DFN2B27. The medium was replaced on day 14. On day 16, after 4 days in static culture, the medium was switched to NBB27 consisting of Neurobasal medium (Gibco) supplemented with 1 × GlutaMAX (Gibco), 1 × NeuroCult SM1 (StemCell Technologies), and 1 × Antibiotic-Antimycotic, and the cultures were then agitated in an incubator shaker (Eppendorf, New Brunswick S41i) at 80 r.p.m. The medium was supplemented with 10 μM SB 431542 (BioGems) from day 0 to day 6, 100 nM LDN-193189 (BioGems) from day 0 to day 12, 100 ng/mL human DKK-1 (PeproTech) and 0.65 μM Purmorphamine (BioGems) from day 6 to day 30, and 20 ng/mL human BDNF (PeproTech), 20 ng/mL human GDNF (PeproTech), 10 ng/mL human IGF-1 (PeproTech) and 100 μM dibutyryl-cAMP (BioGems) from day 30 to day 60. The cells were fed every 3 days from day 0 to day 60 except for day 12 and day 14. From day 60, dibutyryl-cAMP was withdrawn from the medium and the cells were fed every 3-4 days.

### Real-time qPCR

Prior to RNA extraction, organoids were minced with a blade. Total cellular RNA was extracted from small pieces of organoids using TRIzol Reagent (Invitrogen) and a Direct-zol RNA MiniPrep kit (Zymo Research), in accordance with the manufacturer’s instructions. cDNA was prepared by reverse transcription using the SuperScript IV VILO Master Mix (Invitrogen). Real-time qPCR was carried out using QuantiTect SYBR Green PCR Kit (Qiagen) on QuantStudio 3 Real-Time PCR System (Applied Biosystems). The primer sequences are shown in Table S10.

### Immunostaining

Organoids were fixed with 4% PFA in PBS overnight at 4°C. After washing with PBS, organoids were placed in serial dilutions of PBS-buffered sucrose (10, 20, and 30%, in sequence) at 4°C. Each solution was replaced every day. The dehydrated organoids were maintained in 30% sucrose solution at 4°C until embedding with OCT compound (Sakura Finetek). One day before cryosectioning, fixed organoids were placed in a 1:2 mixture of 30% sucrose solution and OCT compound and left overnight at 4°C. Then, organoids were embedded in a 1:2 mixture of 30% sucrose solution and OCT compound, frozen immediately in dry ice/acetone, and cryosectioned at 10 μm. Tissue sections were subjected to additional fixation with 4% PFA in PBS for 3 min at room temperature, following which they were permeabilized and blocked with 10% normal donkey serum in PBS containing 0.3% Triton X-100 for 30 min at room temperature. After washing with 5% serum in PBS containing 0.01% Tween-20, sections were incubated with primary antibodies at 4°C overnight and with secondary antibodies at room temperature for 90 min. Detailed information regarding antibodies is available in Table S11.

### Multi-electrode array (MEA) recording

VFOs were plated per well in 48-well MEA plates (Axion Biosystems) on day 130. The plate was coated with 0.1% Polyethylenimine solution (Sigma Aldrich) and 10 μg/mL mouse Laminin (Gibco). Organoids were then fed twice a week with Brain Phys Neuronal medium (StemCell Technologies) supplemented with 1 × NeuroCult SM1, 10 mM glucose, 1 × Antibiotic-Antimycotic, 20 ng/mL human BDNF, 20 ng/mL human GDNF and 10 ng/mL human IGF-1. For the first 7 days, 5 μg/mL of Laminin was added to the medium to enhance the organoid attachment to the plates. The measurements were collected 24 hours after the medium was changed, once a week, starting at 20 days after plating (150 days of organoid differentiation). Recordings were performed using a Maestro MEA system and AxIS Software Spontaneous Neural Configuration (Axion Biosystems). Spikes were detected with AxIS software using an adaptive threshold crossing set to 5.5 times the standard deviation of the estimated noise for each electrode. The plate was first allowed to rest for 3 minutes in the Maestro device, and then data were recorded for 5 minutes. The MEA analysis was performed as previously described [95]. The pharmacological manipulation was performed with VFOs plated on a 48-well MEA plate (n = 3-9, VFO culture) using the following drugs: 10 μM bicuculline, 50 μM muscimol, 20 μM CNQX, 20 μM AP5. In this assessment, baseline recordings were obtained immediately before and 15 minutes after the addition of the compound(s). The organoids were washed three times with the medium and another recoding was conducted after 24 or 48 hours of incubation at 37°C in washout experiments.

### Dissociation of organoids and scRNA-seq

Three to five VFOs were randomly selected from each iPS cell line on days 70 and 150. Organoids were dissociated into single cells by using the Papain dissociation system (Worthington Biochemical) as previously described [96]. To pool multiple samples together into a single library, cell hashing technology [97] was applied, in accordance with the manufacturer’s instructions (BioLegend). Briefly, 1×106 cells from each iPS cell line were resuspended in 45 μL cell staining buffer (BioLegend) in a protein low-binding microcentrifuge tube separately and incubated with 5 μL Fc blocking reagent (BioLegend) for 10 min at 4°C. Pre-centrifuged 50 μL of hashtag antibody solution containing 2 uL of TotalSeq-A Hashtag antibody (BioLegend), with a unique barcode sequence for each iPS cell line, in 48 μL cell staining buffer was added to the 50 μL blocked cell suspension and incubated for 30 min at 4°C. Cells were then washed three times with cell staining buffer and resuspended in 0.4% BSA/PBS per iPS cell line. Hashtagged single cell suspension from multiple iPS cell lines (up to 12 samples) was pooled together and passed through a 35 μm filter (Falcon). Pooled cell concentration was adjusted at a density of 1,000 cells/μL 0.4% BSA/PBS and approximately 16,000 cells were loaded onto a Chromium Single Cell 3’ Chip (10· Genomics) and 3’ gene expression libraries were generated with a Chromium Next GEM Single Cell 3’ Reagent Kits v3.1 (10· Genomics) according to the manufacturer’s instructions. Hashtag oligonucleotide (HTO) libraries were generated in accordance with BioLegend’s instructions (https://www.biolegend.com/en-us/protocols/totalseq-a-antibodies-and-cell-hashing-with-10x-single-cell-3-reagent-kit-v3-3-1-protocol). Libraries were sequenced on an Illumina NovaSeq 6000 (SP, 2·50 bp) by the Genetic Resources Core Facility at Johns Hopkins University, School of Medicine. HTO and 3’ gene expression libraries were sequenced on the same lane with a 1:99 ratio.

### Data analysis for scRNA-seq

Raw sequencing data were preprocessed with Alevin software [98] (v1.1.0), in which reads were aligned to the hg38 human reference genome. The expression data was processed with the toolkit Scanpy [99] (v1.7.2) in Python3. Initially, cells with less than 200 detected genes or with mitochondrial content higher than 20% were excluded. Genes that were not expressed in at least 3 cells were also removed from the analysis. Read counts were normalized by counts per million (CPM) in each cell and subsequently log-transformed. The most variable genes were defined as having an average normalized expression between 0.0125 and 3 and a dispersion greater than 0.5. Then, dimensionality reduction was computed by principal component analysis (PCA) using the selected highly variable genes. The top 100 significant PCs were used to calculate a neighborhood graph using 15 neighbors as local size, which was then embedded in two dimensions using UMAP (Uniform Manifold Approximation and Projection). Cells were then grouped into different clusters by using the Leiden algorithm [100] with a resolution=1.2.

A cluster of mixed neurons both in VFOs was further divided into 3 subclusters consisting of inhibitory-, excitatory- and midbrain dopaminergic (mDA) neurons with the Leiden algorithm based on the expression of specific marker genes for excitatory neurons (*SLC17A6, SLC17A7, TBR1*, and *SATB2*) and mDA neurons (*PITX3, EN1, EN2, NR4A2, DDC, SLC18A2, SLC6A3, LMX1A, LMX1B, FOXA2*, and *MSX1*). Clusters were annotated to specific cell types based on the expression patterns of known marker genes for each cell type (Table S3). Differentially expressed genes (DEGs) for each cell cluster/type compared with all other cell clusters/types were found by ranking genes with the Wilcoxon rank-sum test, and the *p*-values were adjusted for multiple testing using the Benjamini-Hochberg method. Only genes with an adjusted p < 0.05 were considered as differentially expressed.

A comparison of VFO neuronal progenitors and inhibitory neuronal cell (intermediate progenitor cell, IPC and neuron) data to BrainSpan transcriptomic data of microdissected human brain tissue (https://www.brainspan.org/) was performed using VoxHunt R package with default settings. DEGs between CT and SCZ or day 70 and day 150 in each cell type were identified by the Wilcoxon rank-sum test by considering genes that were expressed in more than 20% of cells in each iPS cell line in each cell type (union of genes expressed at least 20% of cells in each iPS cell lines for each cell type). Analyses were controlled for multiple testing with the Benjamini-Hochberg method. An adjusted *p* < 0.05 was defined as statistical significance. Gene Ontology (GO) enrichment and Kyoto Encyclopedia of Genes and Genomes (KEGG) enrichment of DEGs were performed using the g: Profiler package [101]. Enrichment analysis with GWAS significant genes [62] and TWAS significant and differentially expressed genes in postmortem caudate tissues from individuals with SCZ [66] was performed with the Fisher’s exact test.

### Trajectory analysis

RNA velocity analysis [63] on VFO neuronal progenitors (RG, Ventral RG, and oRG), excitatory IPC/neuron, and inhibitory IPC/neurons was performed using the scVelo [64] (v.0.2.4) python package using a dynamical model with differential kinetic analysis. The number of spliced and unspliced reads was counted directly on the Alevin output. Genes with less than 20 counts (for both unspliced and spliced) were filtered out, whereas all DEGs identified in neuronal progenitors, excitatory- and inhibitory IPC/neurons were retained. As a result, 11,292 genes were remaining, of which the 10000 most variable ones were used for next steps. Count normalization, computation of first/second-order moments, velocities estimation and UMAP embedding were all performed with the default parameters of the built-in functions (https://scvelo.readthedocs.io/). Potential driver genes and latent time were also computed with default parameters. Differences in the distribution of each cell type along latent time between CT and SCZ were performed using the Kolmogorov-Smirnov Test. Enrichment of driver genes with DEGs in each cell type of VFOs and SCZ risk genes [62,65,66] were calculated with the Fisher’s exact test.

### Statistics

Data are presented as mean ± s.e.m. or box plots showing maximum, third quartile, median, first quartile, and minimum values. Raw data were tested for normality of distribution, and statistical analyses were performed using unpaired t-tests (two-tailed), one-way ANOVA tests with Tukey’s multiple comparisons, or Kruskal–Wallis tests with Dunn’s multiple comparisons. GraphPad Prism version 7 was used for statistical analyses. *P* values of less than 0.05 were considered to indicate a significant difference between groups.

## ACKNOWLEDGEMENTS

We wish to thank Ms. Bareera Qamar, Dr. Michael McConnell, Dr. Gianluca Ursini and Dr. Danny Chen (Lieber Institute for Brain Development), Mr. David Mohr, Dr. Kakali Sarkar (Johns Hopkins University School of Medicine), Dr. Ann Graybiel (Massachusetts Institute of Technology) and Dr. Anne B. Young and Dr. Cristopher Bragg (Massachusetts General Hospital) for their technical supports, helpful comments and review of this manuscript. This work was supported by the Lieber Institute for Brain Development, the MGH Collaborative Center for X-Linked Dystonia-Parkinsonism (JAE, ACMP), the Maryland Stem Cell Research Fund (LD), and the National Institutes of Health (KJMB - T32MH015330).

## CONFLICT OF INTEREST

The authors declare no conflict of interest.

## Notes

### Competing Interest Statement

The authors have declared no competing interest.

