## Supplementary Figures for "Ventral forebrain organoids derived from individuals with schizophrenia recapitulate perturbed striatal gene expression dynamics of the donor’s brains"

a

| iPS cell line ID | Brain ID | Diagnosis | Race | Sex | PRS | Age of death |
|---|---|---|---|---|---|---|
| LIBD2c1 | Br2371 | Control | Caucasian | Male | - 14.4 | 23 |
| LIBD6c2 | Br5207 | Control | Caucasian | Male | - 10.3 | 62 |
| LIBD7c6 | Br5287 | Control | Caucasian | Male | - 14.7 | 40 |
| LIBD9c1 | Br5212 | Control | Caucasian | Male | - 12.5 | 52 |
| LIBD3c5 | Br5224 | Schizophrenia | Caucasian | Male | - 7.7 | 46 |
| LIBD5c7 | Br5446 | Schizophrenia | Caucasian | Male | - 10.0 | 23 |
| LIBD8c4 | Br2346 | Schizophrenia | Caucasian | Male | - 8.0 | 21 |

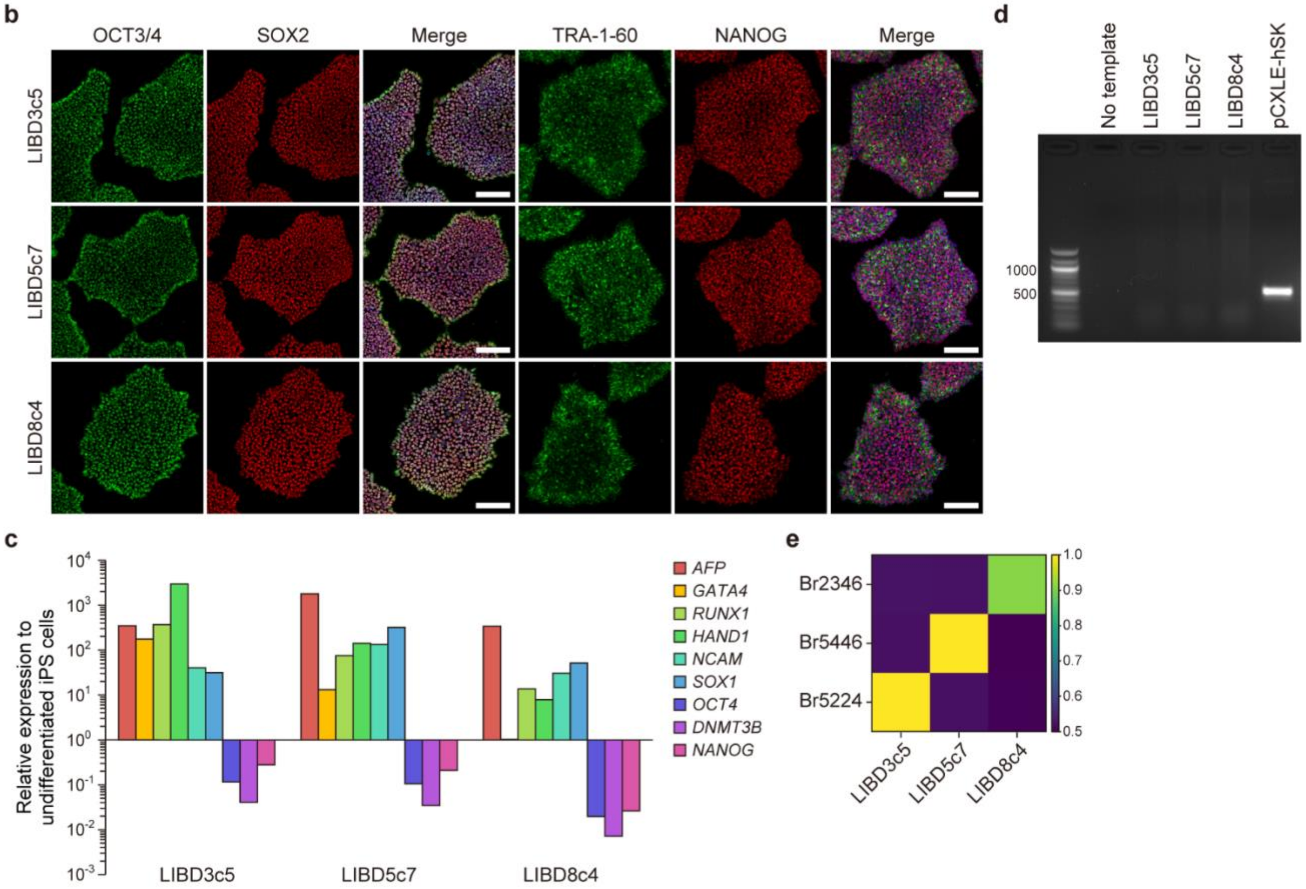


**Supplementary figure 1 | Characterization of postmortem dural fibroblast-derived iPSCs from 3 patients with SCZ.**

**a,** Summary of iPS cell cohort. PRS, polygenic risk score for SCZ.

**b,** Immunostaining of iPSCs derived from postmortem dural fibroblast of 3 individuals with SCZ. Nuclei are stained by Hoechst 33342 (blue) in merged images. Scale bars, 100 µm.

**c**, RT-qPCR result showing expression of marker genes for 3 germ layers (endoderm, *AFP* and *GATA4*; mesoderm, *RUNX1,* and *HAND1*; ectoderm, *NCAM1,* and *SOX1*) and pluripotent stem cells (*OCT4*, *DNMT3B,* and *NANOG*) in the cells induced spontaneous differentiation for 20 days.

**d**, Genomic PCR results showing no genomic integration of episomal vectors used for reprogramming. pCXLE-hSK was used as a positive control.

**e**, Heatmap showing genotype similarity between 3 SCZ iPSC lines and their corresponding postmortem brain tissues.

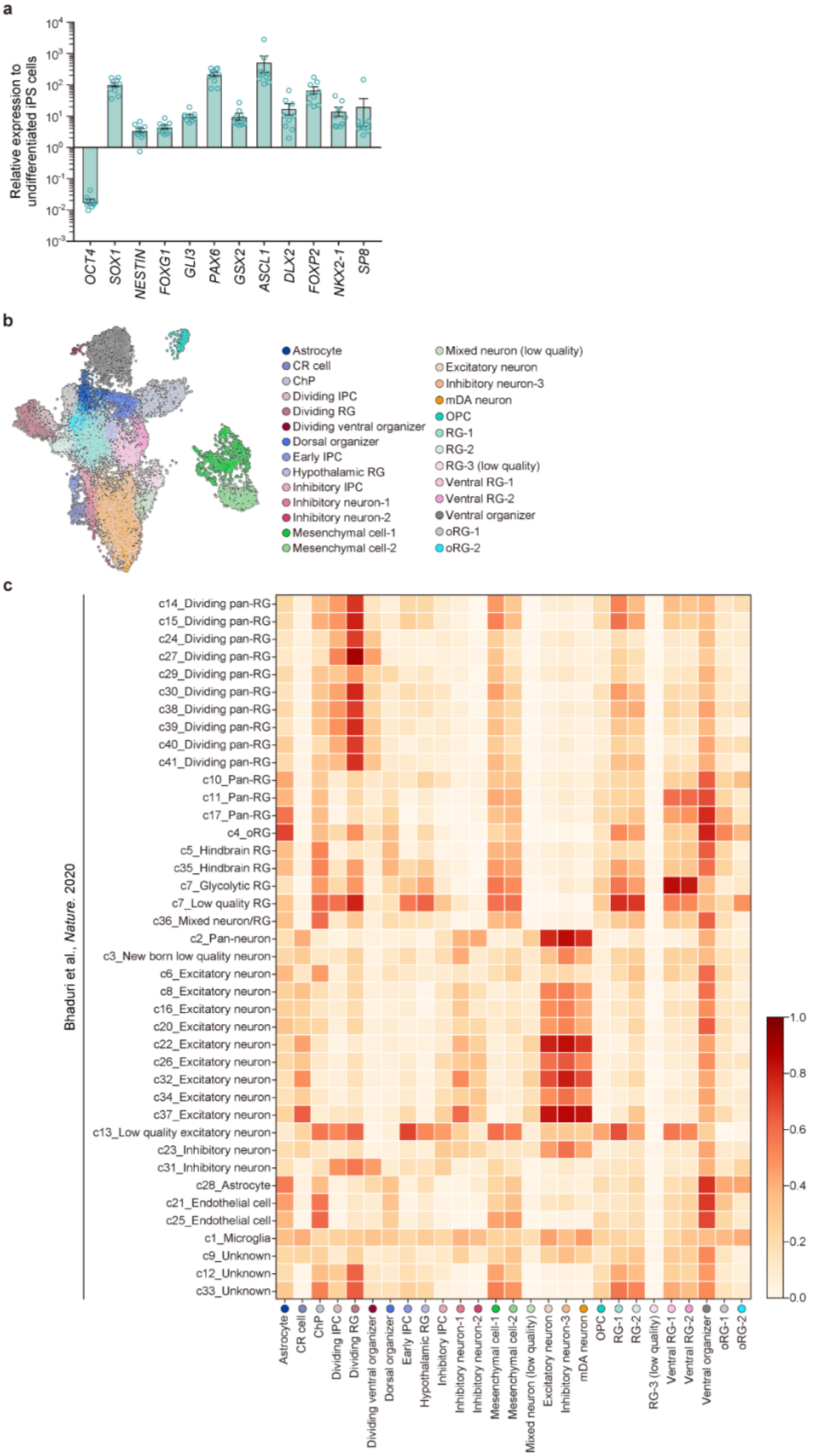
a
Relative expression to undifferentiated iPS cells
OCT4
SOX1
NESTIN
FOXG1
GLI3
PAX6
GSX2
ASCL1
DLX2
FOXP2
NKX2-1
SP8
b
Astrocyte
CR cell
ChP
Dividing IPC
Dividing RG
Dividing ventral organizer
Dorsal organizer
Early IPC
Hypothalamic RG
Inhibitory IPC
Inhibitory neuron-1
Inhibitory neuron-2
Mesenchymal cell-1
Mesenchymal cell-2
Mixed neuron (low quality)
Excitatory neuron
Inhibitory neuron-3
mDA neuron
OPC
RG-1
RG-2
RG-3 (low quality)
Ventral RG-1
Ventral RG-2
Ventral organizer
oRG-1
oRG-2
c
Bhaduri et al., Nature. 2020
c14_Dividing pan-RG
c15_Dividing pan-RG
c24_Dividing pan-RG
c27_Dividing pan-RG
c29_Dividing pan-RG
c30_Dividing pan-RG
c38_Dividing pan-RG
c39_Dividing pan-RG
c40_Dividing pan-RG
c41_Dividing pan-RG
c10_Pan-RG
c11_Pan-RG
c17_Pan-RG
c4_oRG
c5_Hindbrain RG
c35_Hindbrain RG
c7_Glycolytic RG
c7_Low quality RG
c36_Mixed neuron/RG
c2_Pan-neuron
c3_New born low quality neuron
c6_Excitatory neuron
c8_Excitatory neuron
c16_Excitatory neuron
c20_Excitatory neuron
c22_Excitatory neuron
c26_Excitatory neuron
c32_Excitatory neuron
c34_Excitatory neuron
c37_Excitatory neuron
c13_Low quality excitatory neuron
c23_Inhibitory neuron
c31_Inhibitory neuron
c28_Astrocyte
c21_Endothelial cell
c25_Endothelial cell
c1_Microglia
c9_Unknown
c12_Unknown
c33_Unknown
1.0
0.8
0.6
0.4
0.2
0.0

**Supplementary figure 2 | Characterization of ventral forebrain organoids (VFOs) (related to figure 1).**

**a,** RT-qPCR result showing induction of expression of marker genes for neural fate (*SOX1* and *NESTIN*), forebrain (*FOXG1*), dorsal forebrain (*GLI3* and *PAX6*), ventral forebrain (*GSX2*, *ASCL1,* and *DLX2*), LGE (*FOXP2*), MGE (*NKX2-1*) and CGE (*SP8*) and suppression of pluripotent stem cell marker (*OCT4*) in VFOs from 3 CT (LIBD2c1, LIBD7c6, and LIBD9c1) on day 12 (n = 3 for each iPSC line; error bars, s.e.m.).

**b**, UMAP showing 15,424 cells from VFOs of 4 CT and 3 SCZ on days 70 and 150, colored by 26 cell clusters.

**c**, Heatmap showing overlap of DEGs in each of 26 cell clusters compared to all other clusters in VFOs with marker genes for 41 cell clusters in cerebral organoids [102]. The color indicates the overlap coefficient.

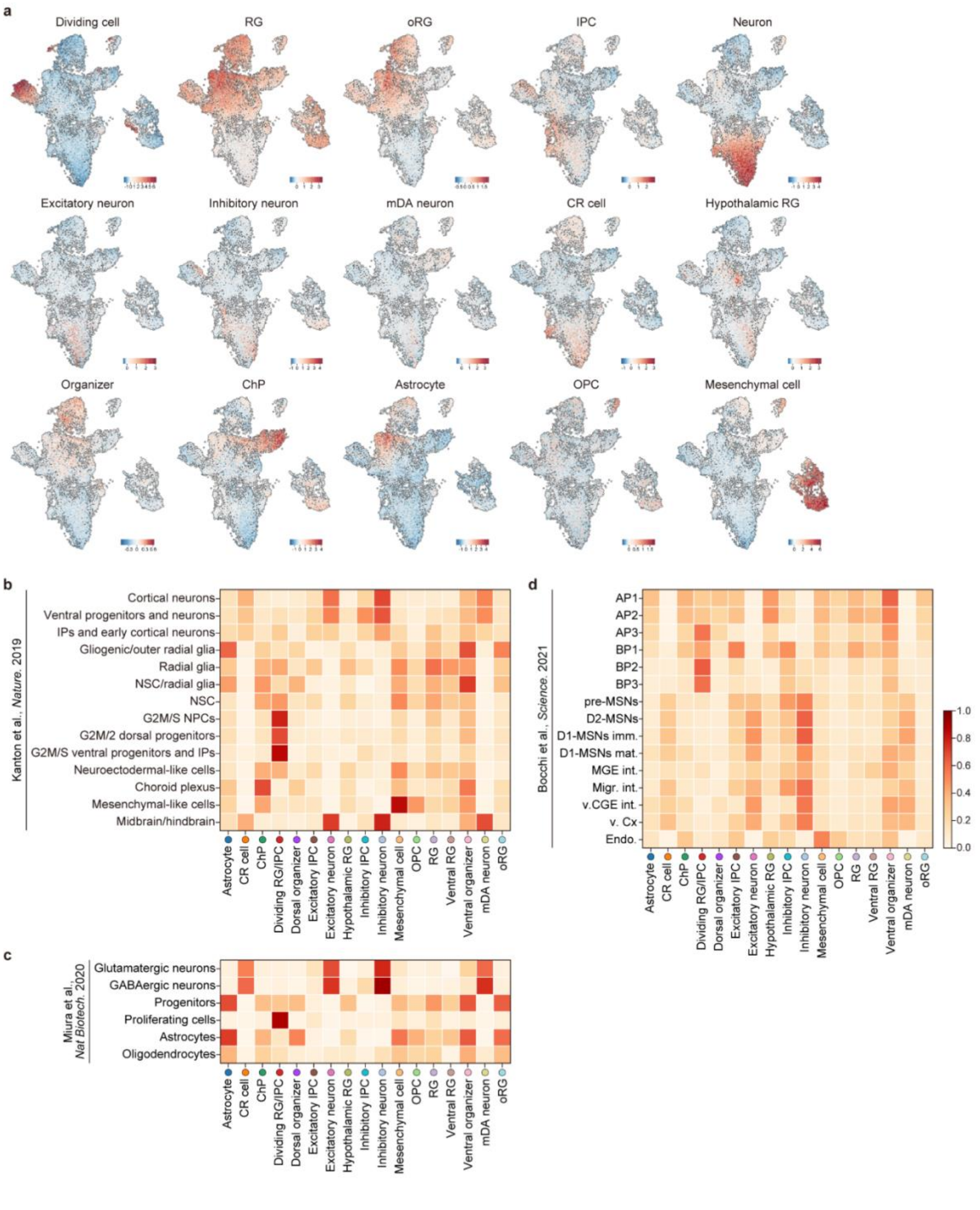


**Supplementary figure 3 | Cell type annotation of VFOs (related to figure 1).**

**a,** Representative mean expression of selected marker genes for each cell type visualized by UMAP. **b**-**d**, Heatmap showing overlap of DEGs in each of 17 cell clusters compared to all other cell types excluding 'Outlier' in VFOs with marker genes for 14 cell types in cerebral organoids [38] (**b**), 6 cell types in striatal organoids [39] (**c**) and 15 cell types in human LGE at PCW 7, 9 and 11 [40] (**d**). Color indicates overlap coefficient.

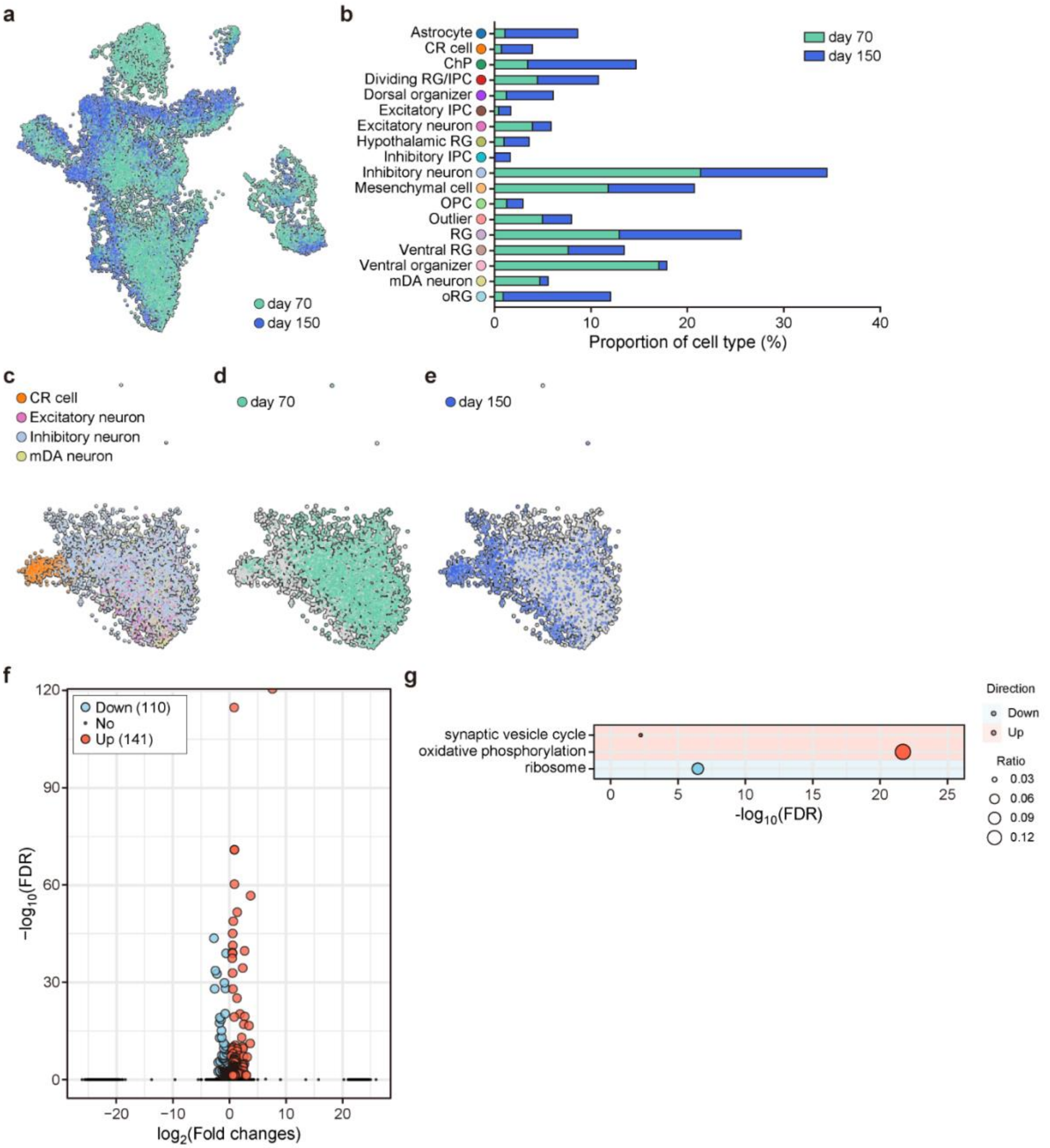


**Extended figure 4 | Comparison of VFOs between day 70 and day 150.**
**a**, UMAP showing 15,424 cells from VFOs of 4 CT and 3 SCZ, colored by day.
**b**, Proportion of 18 major cell types in VFOs on day 70 and day 150.
**c**, UMAP of neurons in VFOs (4,050 cells) from 4 CT and 3 SCZ, colored by cell type.
**d**-**e**, UMAP of neurons. Cells on day 70 (2,828 cells, **d**) and on day 150 (1,222 cells, **e**) are colored on the UMAP.
**f**, Volcano plot showing DEGs in VOFs on day 150 compared with VFOs on day 70.
**g**, Dot plot showing significantly enriched KEGG pathways for upregulated and downregulated genes on day 150 compared to VFOs on day 70. Size of circles indicates ratio of genes annotated in each pathways to all 38,866 genes analyzed. DFR, false discovery rate.

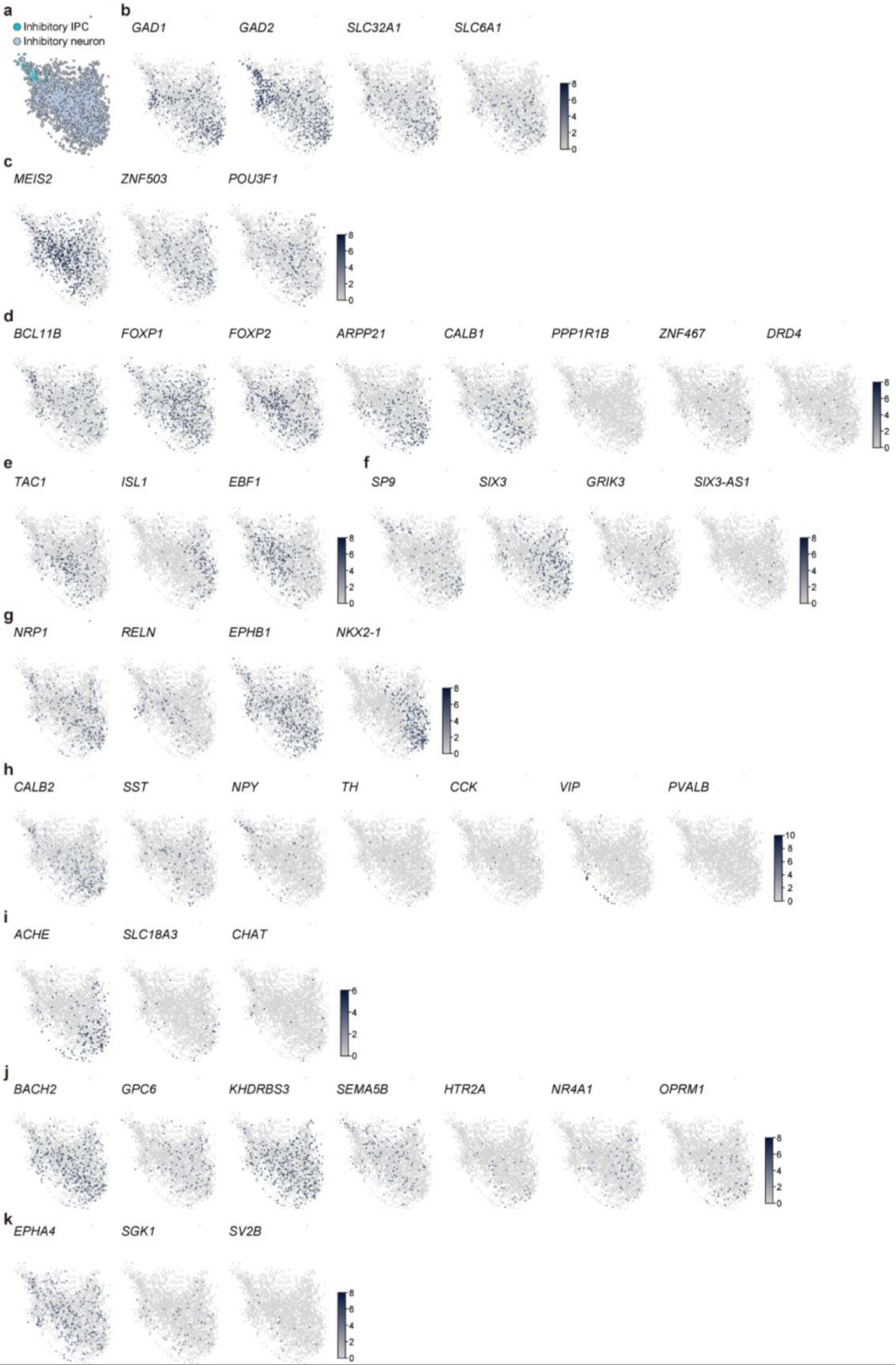
a
Inhibitory IPC
Inhibitory neuron
b
GAD1
GAD2
SLC32A1
SLC6A1
c
MEIS2
ZNF503
POU3F1
d
BCL11B
FOXP1
FOXP2
ARPP21
CALB1
PPP1R1B
ZNF467
DRD4
e
TAC1
ISL1
EBF1
f
SP9
SIX3
GRIK3
SIX3-AS1
g
NRP1
RELN
EPHB1
NKX2-1
h
CALB2
SST
NPY
TH
CCK
VIP
PVALB
i
ACHE
SLC18A3
CHAT
j
BACH2
GPC6
KHDRBS3
SEMA5B
HTR2A
NR4A1
OPRM1
k
EPHA4
SGK1
SV2B

**Supplementary figure 5 | Characterization of inhibitory neuronal cells in VFOs (related to figure 2).**

**a,** UMAP of inhibitory neuronal population in VFOs (2,896 cells from 4 CT and 3 SCZ).

**b**, Expression of representative marker genes for GABAergic neurons visualized by UMAP of inhibitory IPC/neurons.

**c**-**d**, Expression of representative marker genes for pre-medium spiny neurons (MSN) (**c**) and pan-MSN (**d**) visualized by UMAP of inhibitory IPC/neurons.

**e**-**f**, Expression of representative marker genes for pre-D1 MSN (**e**) and pre-D2 MSN (**f**) visualized by UMAP of inhibitory IPC/neurons.

**g**, Expression of representative marker genes for cortical interneurons (*NRP1* and *RELN*) and striatal interneurons (*EPHB1* and *NKX2-1*) visualized by UMAP of inhibitory IPC/neurons.

**h**-**i**, Expression of representative marker genes for striatal GABAergic interneurons (**h**) and cholinergic interneurons (**i**) visualized by UMAP of inhibitory IPC/neurons.

**j**-**k**, Expression of representative marker genes for striosome (**j**) and matrix (**k**) visualized by UMAP of inhibitory IPC/neurons. Expression levels are shown in $\log_2$(cpm).

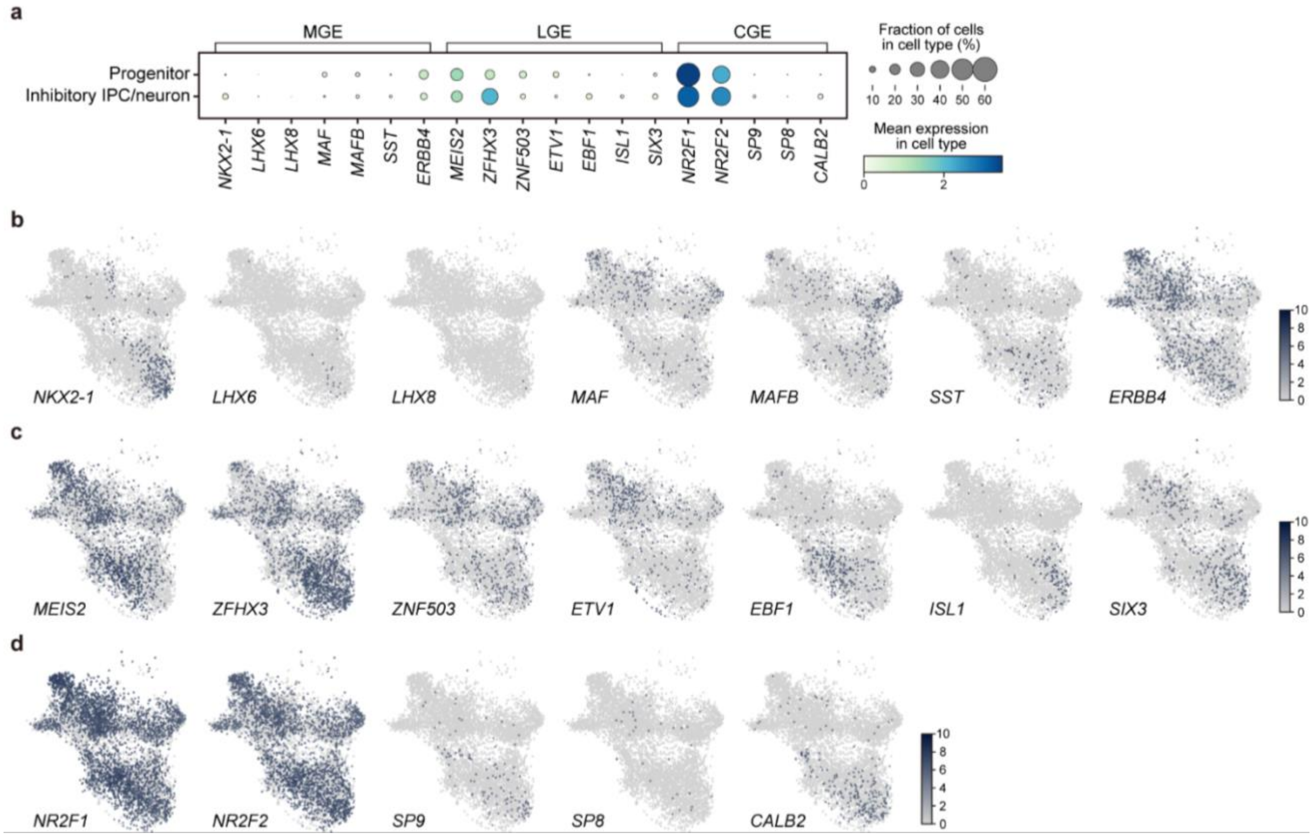


**Supplementary figure 6 | Characterization of neuronal progenitors and inhibitory neuronal cells in VFOs (related to figure 2).**

**a,** Dot plot showing expression of selected marker genes for medial ganglionic eminence (MGE) lateral ganglionic eminence (LGE), and caudal ganglionic eminence (CGE) and percentage of cells expressing those markers in neuronal progenitors and inhibitory neuronal population.

**b**-**d**, Expression of selected marker genes for MGE (**b**), LGE (**c**), and CGE (**d**) visualized by UMAP of neuronal progenitors and inhibitory IPC/neuron in VFO (6,747 cells) from 4 CT and 3 SCZ. Expression levels are shown in $log_2$(cpm).

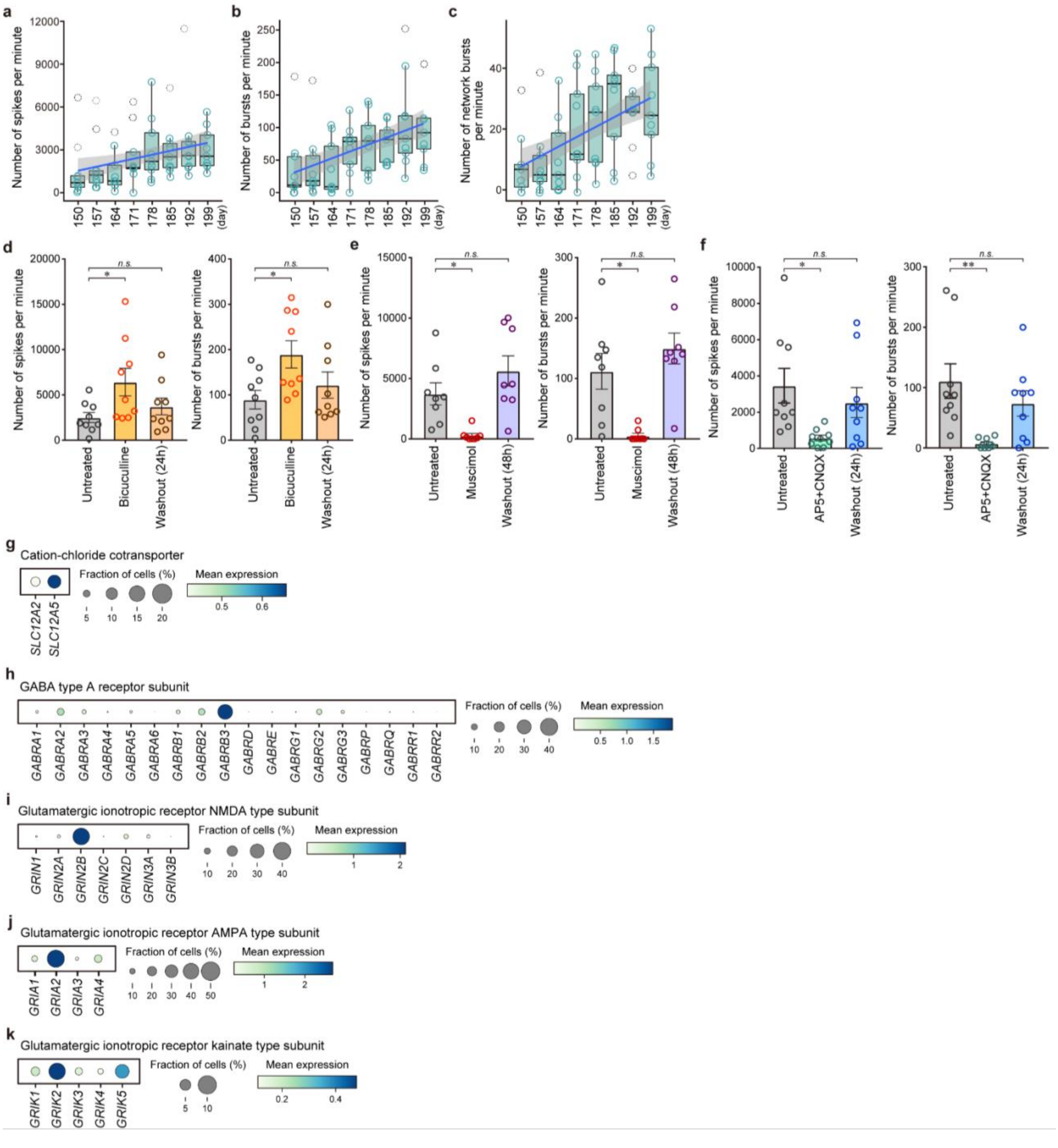

a
Number of spikes per minute
150 157 164 171 178 185 192 199 (day)
b
Number of bursts per minute
c
Number of network bursts per minute
d
Untreated
Bicuculline
Washout (24h)
n.s.
*
Number of bursts per minute
e
Muscimol
Washout (48h)
f
AP5+CNQX
**
g
Cation-chloride cotransporter
SLC12A2
SLC12A5
Fraction of cells (%)
Mean expression
h
GABA type A receptor subunit
GABRA1 GABRA2 GABRA3 GABRA4 GABRA5 GABRA6 GABRB1 GABRB2 GABRB3 GABRD GABRE GABRG1 GABRG2 GABRG3 GABRP GABRQ GABRR1 GABRR2
i
Glutamatergic ionotropic receptor NMDA type subunit
GRIN1 GRIN2A GRIN2B GRIN2C GRIN2D GRIN3A GRIN3B
j
Glutamatergic ionotropic receptor AMPA type subunit
GRIA1 GRIA2 GRIA3 GRIA4
k
Glutamatergic ionotropic receptor kainate type subunit
GRIK1 GRIK2 GRIK3 GRIK4 GRIK5

**Supplementary figure 7 | Functional characterization of VFOs (related to figure 2).**
**a**-**c**, Time series of the number of spikes (**a**), number of bursts (**b**), and number of network bursts (**c**) per minute of VFOs derived from a control individual (33114.c) between day 150 and day 199 (n = 3-9; VOFs for each time point). Dashed circles indicate outliers.
**d**-**f**, Effect of treatments with bicuculline (**d**), muscimol (**e**) and AP5+CNQX (**f**) on the number of spikes and number of bursts per minute of control VFOs at day 220 (**d**), day 241 (**e**), and day 227 (**f**). Data represent mean ± s.e.m. (n = 9; VOFs for recording; one-way ANOVA; $^{*}p<0.05$; $^{**}p<0.01$; n.s., not significant).
**g**, Dot plot showing expression of cation-chloride transporter (**g**) and percentage of cells expressing those genes in inhibitory neurons in VFOs from 4 CT and 3 SCZ.
**h-k**, Dot plot showing expression of GABA type A receptor subunit (**h**), NMDA type subunit (**i**), AMPA type subunit (**j**), and kainate type subunit (**k**) of glutamatergic ionotropic receptor and percentage of cells expressing those genes in neurons in VFOs from 4 CT and 3 SCZ.

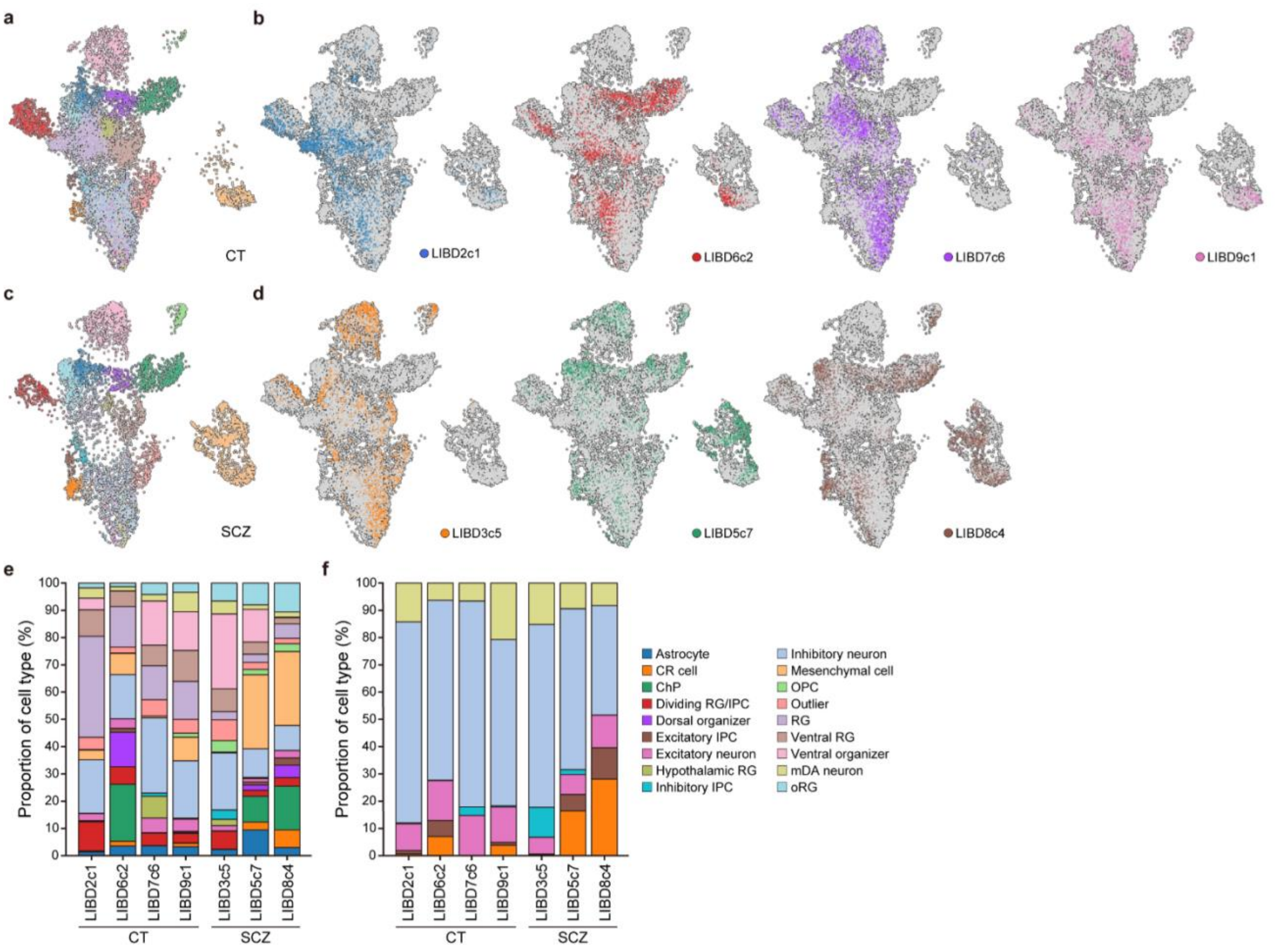


**Supplementary figure 8 | Variation in cell type proportion of VFOs among iPSC lines.**

**a**, UMAP of VFOs derived from 4 control individuals, colored by 18 major cell types.

**b**, Cells belonging to each of 4 CT iPSC lines are visualized individually on the UMAP of VFOs from 7 iPSC lines (4 CT and 3 SCZ).

**c**, UMAP of VFOs derived from 3 individuals with SCZ, colored by 18 major cell types.

**d**, Cells belonging to each of 3 SCZ iPSC lines are visualized individually on the UMAP of VFOs from 7 iPSC lines (4 CT and 3 SCZ).

**e**-**f**, Proportion of 18 major cell types (**e**) and 6 major cell types among IPC/neuron population (**f**) in VFOs from each iPSC line.

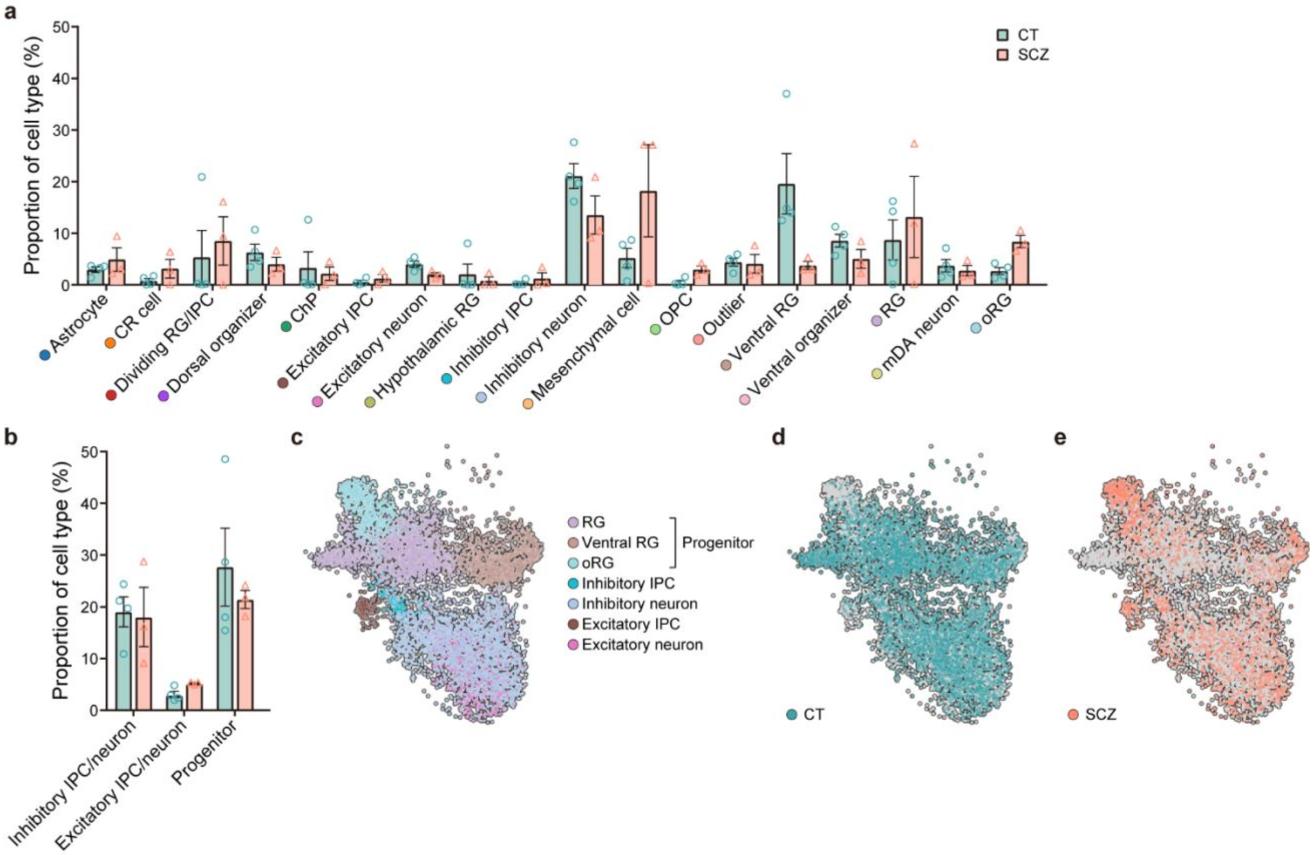


**Supplementary figure 9 | Differences in cell type proportion of VFOs between CT and SCZ.**

**a**, Proportion of each of 18 major cell types in CT VFOs and SCZ VFOs. Data represent mean ± s.e.m.

**b**, Proportion of neuronal cell types in CT VFOs and SCZ VFOs. Data represent mean ± s.e.m.

**c**, UMAP of neuronal progenitors and excitatory and inhibitory neuronal population in VFOs (15,424 cells) from 4 CT and 3 SCZ, colored by major cell type.

**d**-**e**, Cells belonging to CT (8,297 cells from 4 lines, **d**) and SCZ (7,127 cells from 3 lines, **e**) are colored on the UMAP of neuronal progenitors and excitatory and inhibitory neuronal population.

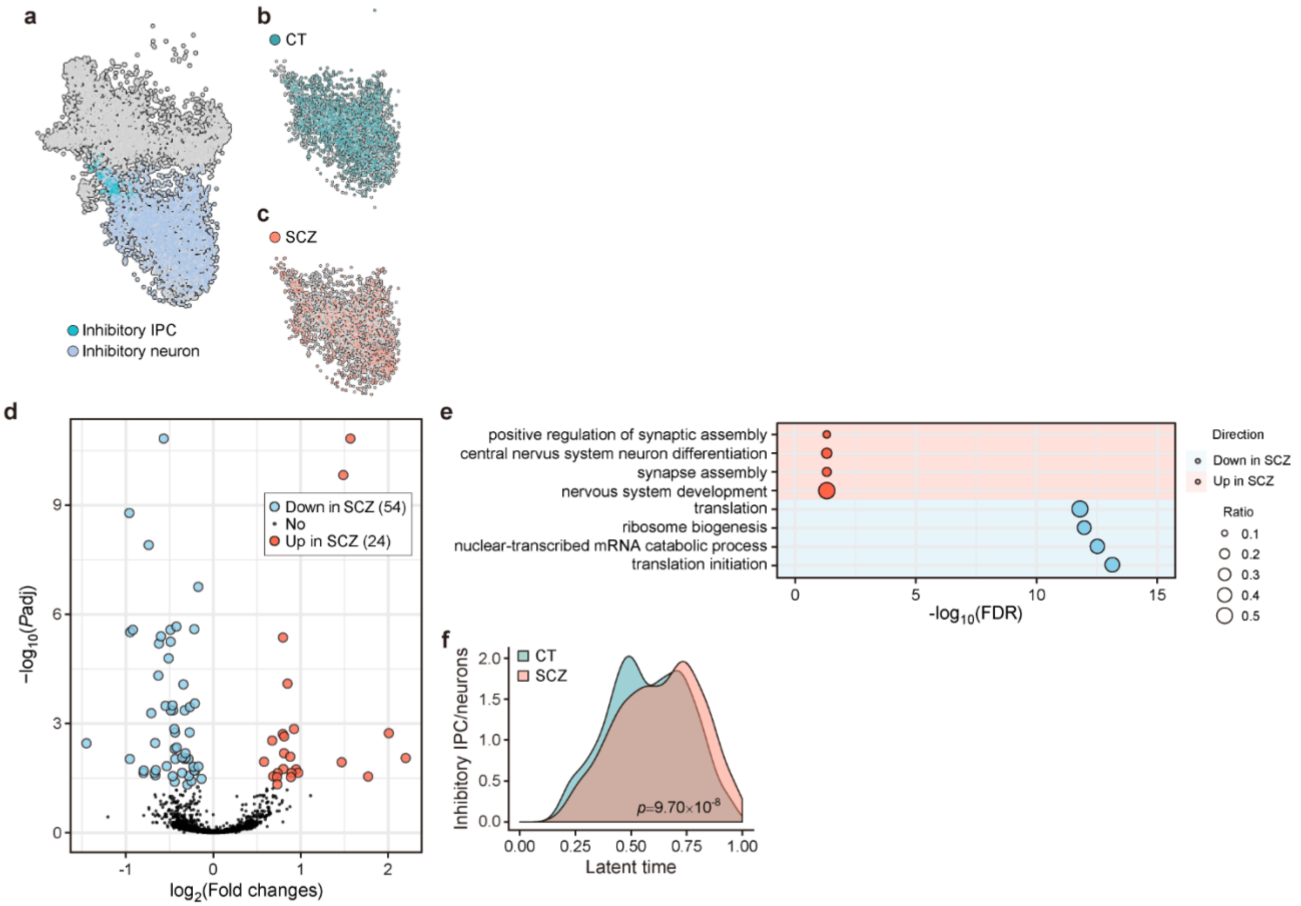


**Supplementary figure 10 | Inhibitory IPC/neuron-specific differences in a gene expression profile and neurodevelopmental trajectory between CT and SCZ.**

**a**, Cells belonging to inhibitory IPC/neurons are colored on the UMAP of neuronal progenitors and excitatory and inhibitory IPC/neurons in VFOs from 4 CT and 3 SCZ.

**b**-**c**, Cells belonging to CT (1,568 cells from 4 lines, **b**) and SCZ (1,328 cells from 3 lines, **c**) are colored on the UMAP of inhibitory IPC/neurons.

**d**, Volcano plot showing DEGs in SCZ compared with CT. Padj, adjusted *p*-value.

**e**, Dot plot showing representative GO terms enriched for upregulated and downregulated genes in SCZ inhibitory IPC/neurons. Size of circles indicates ratio of genes annotated in each GO term to all 2,204 genes analyzed. FDR, false discovery rate.

**f**, Difference in the density of inhibitory IPC/neurons along latent time (Kolmogorov-Smirnov test).

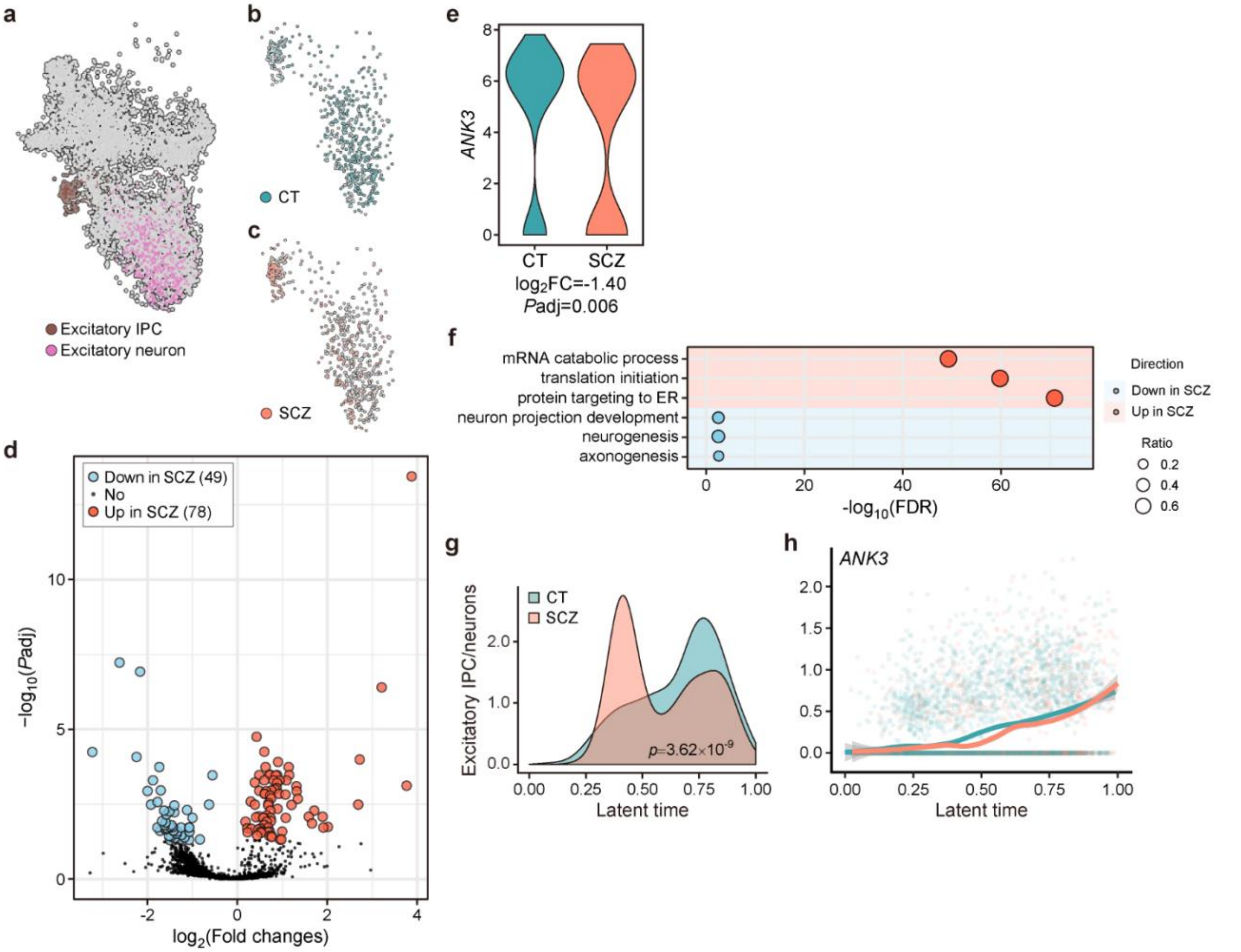


**Supplementary figure 11 | Excitatory IPC/neuron-specific differences in a gene expression profile and neurodevelopmental trajectory between CT and SCZ.**

**a**, Cells belonging to excitatory IPC/neurons are colored on the UMAP of neuronal progenitors and excitatory and inhibitory IPC/neurons in VFOs from 4 CT and 3 SCZ.

**b**-**c**, Cells belonging to CT (250 cells from 4 lines, **b**) and SCZ (375 cells from 3 lines, **c**) are colored on the UMAP of excitatory IPC/neurons.

**d**, Volcano plot showing DEGs in SCZ compared with CT. Padj, adjusted *p*-value.

**e**, Representative expression of downregulated gene *ANK3* in SCZ excitatory IPC/neurons. FC, fold change.

**f**, Dot plot showing representative GO terms enriched for upregulated and downregulated genes in SCZ Excitatory IPC/neurons. Size of circles indicates ratio of genes annotated in each GO term to all 3,370 genes analyzed. FDR, false discovery rate.

**g**, Difference in the density of excitatory IPC/neurons along latent time (Kolmogorov-Smirnov test).

**h**, Expression of *ANK3*, a driver gene for excitatory IPC/neurons that was downregulated in SCZ along latent time in CT and SCZ VFOs.

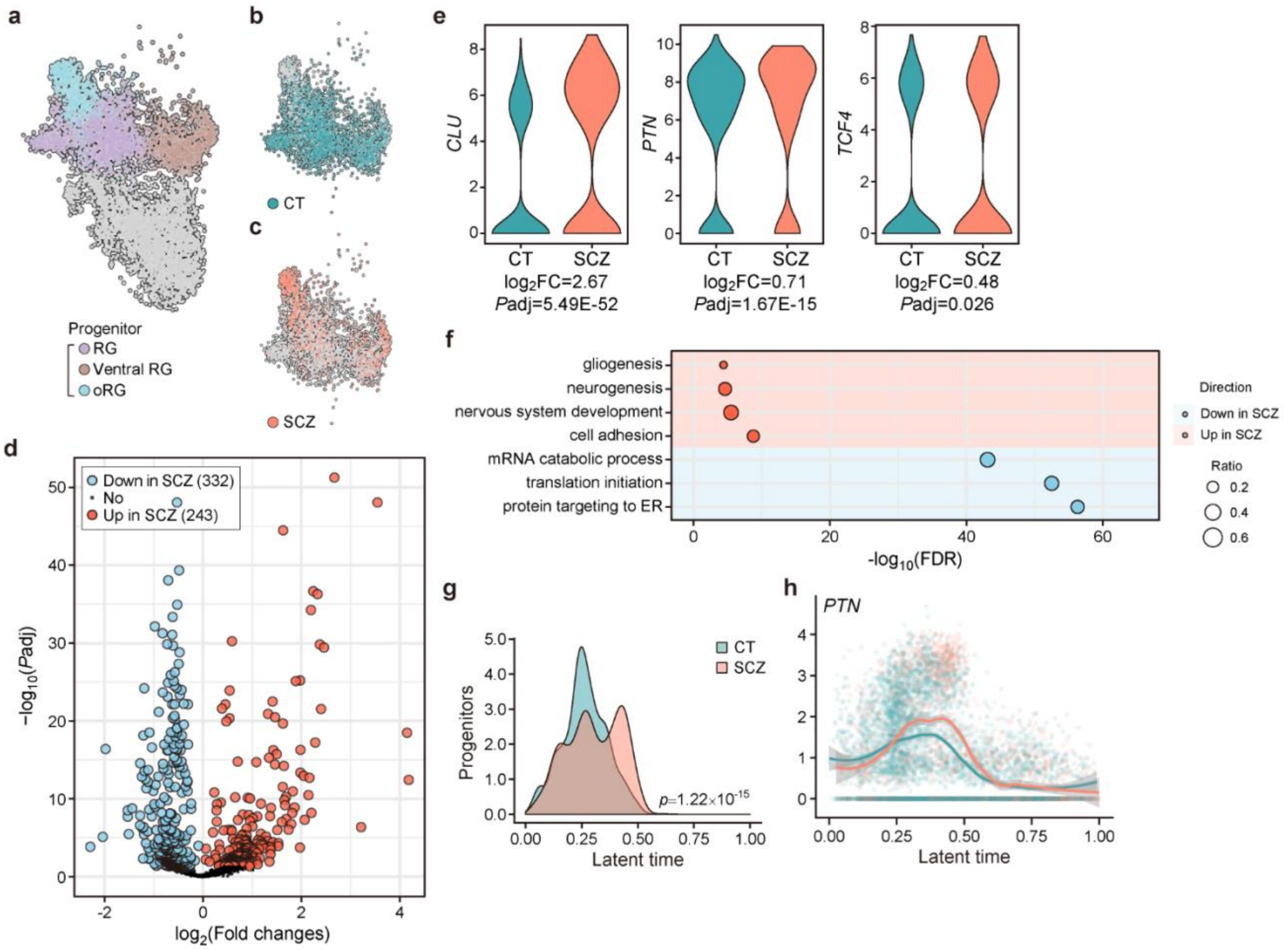


**Supplementary figure 12 | Neuronal progenitor-specific differences in a gene expression profile and neurodevelopmental trajectory between CT and SCZ.**

**a**, Cells belonging to neuronal progenitors are colored on the UMAP of neuronal progenitors and excitatory and inhibitory IPC/neurons in VFOs from 4 CT and 3 SCZ.

**b**-**c**, Cells belonging to CT (2,315 cells from 4 lines, **b**) and SCZ (1,536 cells from 3 lines, **c**) are colored on the UMAP of neuronal progenitors.

**d**, Volcano plot showing DEGs in SCZ compared with CT. Padj, adjusted *p*-value.

**e**, Representative expression of GWAS-significant genes upregulated in SCZ progenitors. FC, fold change.

**f**, Dot plot showing representative GO terms enriched for upregulated and downregulated genes in SCZ progenitors. Size of circles indicates ratio of genes annotated in each GO term to all 2,380 genes analyzed. FDR, false discovery rate.

**g**, Difference in the density of progenitors along latent time (Kolmogorov-Smirnov test).

**h**, Expression of *PTN*, a driver gene for progenitors that was upregulated in SCZ along latent time in CT and SCZ VFOs.

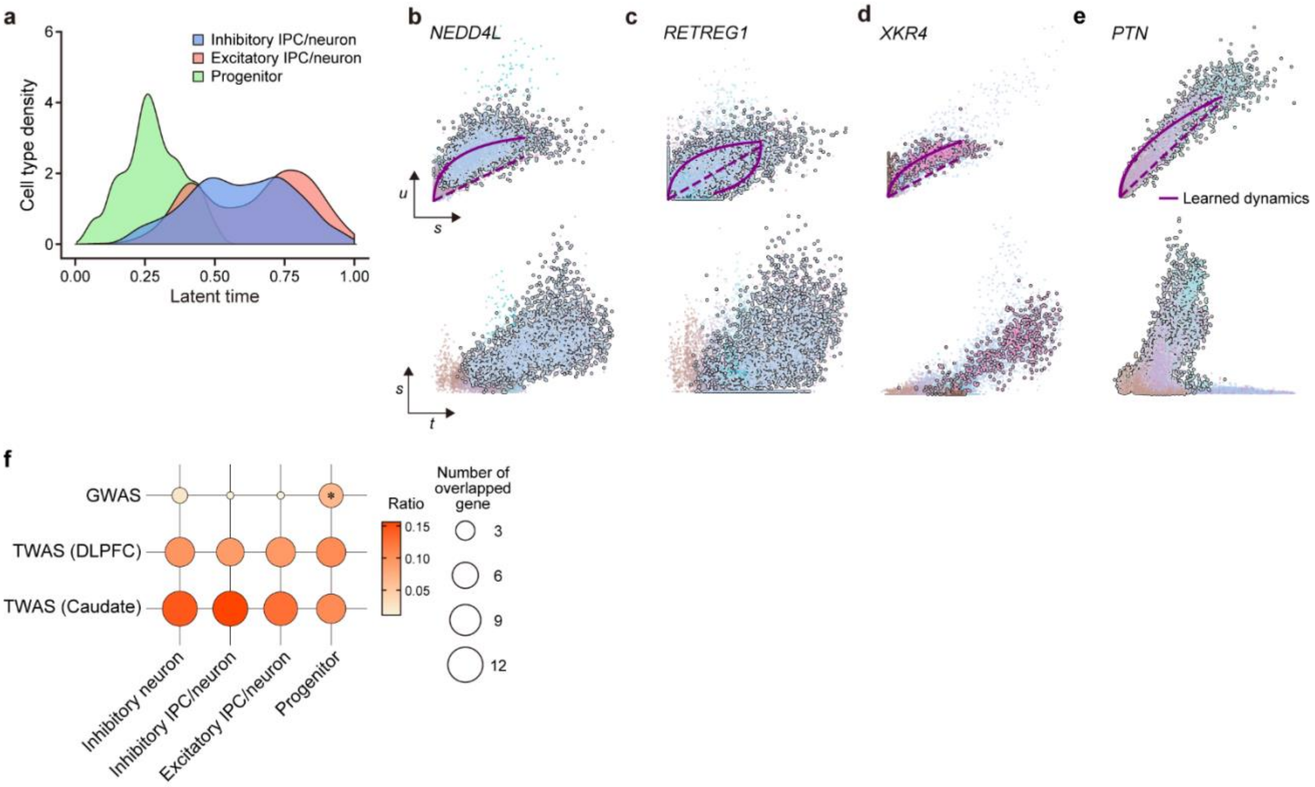


**Supplementary figure 13 | Neurodevelopmental trajectory and driver genes in VFOs.**

**a**, Density of pooled cell types along latent time.

**b**-**e**, Phase portraits (top) and expression dynamics along latent time (bottom) for putative driver genes specific for inhibitory neurons (*NEDDL4*, **b**), inhibitory IPC/neurons (*RETREG1*, **c**), excitatory IPC/neurons (*XKR4*, **d**) and neuronal progenitors (*PTN*, **e**). *u*, unspliced mRNA; *s*, spliced mRNA; *t*, latent time.

**f**, Dot plot showing the overlap between putative drivers for each cell type and SCZ GWAS significant genes [62], SCZ TWAS significant genes in DLPFC [65], and SCZ TWAS significant genes in caudate [66]. Fisher's exact test; $*p<0.05$. The ratio of drivers overlapped with SCZ risk genes to all drivers in each cell type is indicated by color.

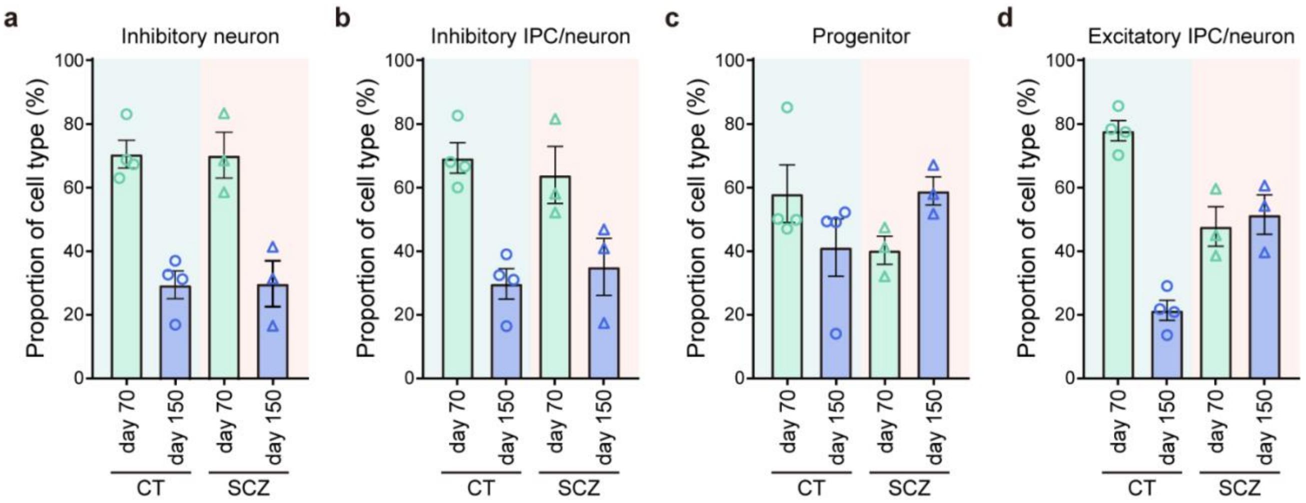


**Supplementary figure 14 | Proportion of neuronal cells from VFOs on day 70 and day 150.**
**a**-**d**, Proportion of inhibitory neurons (**a**), inhibitory IPC/neurons (**b**), neuronal progenitors (**c**), and excitatory IPC/neurons (**d**) in VFOs from CT and SCZ, plotted by day. Data represent mean ± s.e.m.
